# Evolutionary implications of plant host interactions with a generalist pathogen

**DOI:** 10.1101/2025.05.05.652313

**Authors:** Hanna Märkle, Theo L. Gibbs, Joy Bergelson

## Abstract

It is widely believed that eco-evolutionary feedbacks arising from host-pathogen interactions shape the number and frequency of resistance (*R*) genotypes and their allelic polymorphism. A subset of *R* genes exhibit unusually strong signatures of balancing selection, sometimes even existing as a trans-specific polymorphism. Here, we explore the role of alternative hosts on *R* gene evolution through a simple model of two closely related host species that share a single generalist pathogen. We ask (i) how shared interactions determine the *R* gene repertoires and polymorphism in each host and (ii) under which circumstances (trans-specific) polymorphism is maintained. Our results indicate that interactions with a generalist pathogen are more likely to sustain polymorphism at a shared *R* gene compared to the maintenance of polymorphism at *R* genes private to each host. The former can translate into trans-specific resistance gene polymorphism. Further, we observe that increasing the relative proportion of a single host species favors fixation of private resistance in the less common host, which stays effective because the pathogen tracks the more common host. Our model thus sheds light on how *R* gene dynamics are shaped by interactions with a shared generalist pathogen and the pathogen’s response to alternative hosts.

## 1. Introduction

Interactions between hosts and pathogens are widespread in nature. They affect not only the short-term population (ecological) dynamics of the interacting species but also the long-term evolution of genes at their interface, with important implications for agriculture and medicine. This has led to a broad interest in the ecological and population genetic dynamics of host-pathogen systems, beginning with the seminal work of Anderson and May 1982 (1; 2) and continuing to the current day (3; 4). Epidemiological dynamics can alter host and pathogen population sizes, and allele frequencies at the underlying genes change as a consequence of the reciprocal selective pressures that interacting partners impose upon each other. This is especially pronounced when the respective host (pathogen) phenotypes vary greatly in their resistance (infection) success against different genotypes and adhere to fitness trade-offs between growth and resistance (infection).

One biological system conforming to these interactions involves plant resistance *R* genes that play a crucial role in effector-triggered immunity, a key component of the plant immune system (5). These *R* genes recognize small proteins, known as effectors, that bacterial and fungal pathogens secrete into the plant to interfere with basal plant defense responses, thereby promoting their own growth. Interactions between host resistance genes and pathogen effector genes are often characterized by a gene-for-gene (GFG) interaction model (6; 7). Here, successful recognition of a pathogen effector protein by the corresponding host R protein triggers resistance, whereas susceptibility results from the lack of a cognizant host allele or from the lack of a recognized effector gene.

Simple host-pathogen interaction models (one locus with two alleles, in each of the host and pathogen) have demonstrated a role of (in)direct negative-frequency dependent selection driving allelic changes in both interacting partners (8; 9). The frequency of a resistance allele increases when rare, as it provides a fitness advantage compared to susceptible hosts in interactions with pathogens with the recognized effector (avirulent pathogens). This in turn favors an increase in the frequency of virulent pathogen genotypes that escape recognition by the specific resistance allele either due to a mutation in the effector or the loss of the effector. If resistance comes at a genotypic cost, the frequency of susceptiblity will increase because now both resistant and susceptible hosts are equally infected, making the cost of resistance the main selective force. As the frequency of susceptibility increases, there should be an associated increase in the frequency of avirulent pathogens if there is a genotypic cost for virulent pathogens (for a summary see (9)). Thus, allele frequencies fluctuate as result of the specific trade-offs between the costs and benefits of resistance in the host and virulence in the pathogen. In general, the presence of genotypic costs of host resistance and pathogen virulence are necessary but not sufficient conditions for maintaining polymorphism in simple host-pathogen interaction models (9).

As described above, genotypic costs are key components of the dynamics leading to long-lived polymorphisms. There is mixed evidence for fitness costs associated with resistance and virulence (10; 11; 12; 13). For example, the cost of the resistance allele *Rps5* (14) in *Arabidopsis thaliana* has been estimated to reduce the lifetime fitness of resistant genotypes by 5%-10% relative to susceptible genotypes when grown in the absence of the pathogen, but no such differential fitness was detected between pairs of resistant and susceptible lines for the resistance gene *Rps2* (15). Similarly, pathogens lacking a given effector may experience fitness costs to varying extents; these costs manifest themselves as decreased virulence and growth within susceptible plants relative to strains with a functional effector allele (11; 12).

Depending on the specific shape of the genotypic fitness trade-offs, life-history traits of the interacting species and the strength of negative frequency dependent selection, alternative host-pathogen dynamics can result. Under arms-race dynamics (16) resistance alleles/genes and effector alleles/genes that escape recognition are recurrently fixed. This contrasts with “trench-warfare dynamics” (17), where resistance and effector polymorphisms are maintained as a consequence of alternative alleles being favored at different points in time and space. Analyses of data from host-pathogen systems have revealed evidence for both arms races and trench warfare dynamics in natural systems, agriculture and humans (14; 17; 18; 19; 20; 21; 22; 23; 24; 25; 26). In some cases, these resistance polymorphisms have been shown to predate the split of two closely related species and exist as trans-specific polymorphisms (26; 27; 28; 29; 30).

Several factors, such as asynchrony between host and pathogen life-cycles and epidemiological feedbacks, promote polymorphism (9) An integrative study of the long-term stable maintenance of the presence-absence polymorphism at the *Rps5* resistance gene in *A. thaliana* concluded that this ancient balanced polymorphism is best explained by diffuse interactions involving several host species and pathogens with multiple non-homologous effectors (14). Diffuse interactions are likely to be common because a large fraction of pathogen species interact with multiple host species, especially those that are closely related (31; 32) and thus, diffuse interactions between hosts and pathogens may be a common selective agent to promote trans-specific polymorphism.

Community-wide interactions more broadly may play a key role in shaping host-pathogen interactions and coevolutionary dynamics. Work in grassland communities, for example, has highlighted that the disease pressure on individual species was not determined by a species’ relative abundance alone, but also by the abundance of closely related host species, presumably because they can serve as alternative hosts for generalist pathogens (33). Another system implicating community-wide dynamics in host-pathogen interactions is the apparent maladaptation of strains of the generalist pathogen *Pseudomonas syringae* to local genotypes of its host, *A. thaliana*. Here, host genotypes were more likely to recognize local isolates (34), which runs counter to the common assumption that pathogens would be ahead in an arms race (35; 36). This maladaptation has been hypothesized to result from generalist pathogens being more strongly selected to adapt to highly abundant and long-lived hosts, rather than to less abundant and short-lived hosts such *A. thaliana*. This may have allowed for an increase in the frequency of resistance in *A thaliana*, without counter-adaptation in the pathogen to escape recognition.

Given the ubiquity and importance of generalist pathogens, we asked whether interactions between a generalist pathogen and two closely related host species that share some, but not all resistance genes, could maintain *R* gene polymorphisms. To address this question, we constructed a multi-locus gene-for-gene model incorporating two phylogenetically related hosts interacting with a single generalist pathogen. Our intuition was that shared resistance imposes a common selective pressure for the pathogen, while private resistance loci generate host species-specific pressures. Thus, we expected that the relative proportion of the two host species would play a crucial role in determining which genotypes are maintained. With equal frequencies of both host species, we expected frequent maintenance of polymorphism at the shared and the private resistance loci, with the costs of resistance and virulence functions driving the specific outcomes. As one host becomes more common, this balance should shift as any given pathogen genotype is expected to more frequently interact and evolutionarily respond to the more common host. Because encounters with the private resistance allele in the less common host become more rare, and pathogens need to retain some effectors (Avirulence) alleles to be pathogenic, we expect that this situation often leads to an increase in frequency of the less common host’s resistance allele and the matching Avirulence allele in the pathogen.

## 2. Methods and Material

### (a) A two host-generalist pathogen gene-for-gene (GFG) model

We extended existing multi-locus gene-for-gene (GFG) models (8; 37) to incorporate two phylogenetically related host species *H* and *M* interacting with a single generalist pathogen *P*. Hosts have relative proportions *ϕ* (for host *H*) and (1 − *ϕ*) (for host *M*). We assume that (i) there is no competition between the two host species *H* and *M*, and (ii) there is no preference of the pathogen for either host, beyond certain pathogen genotypes being better at infecting certain host genotypes based on their *R* gene repertoires.

#### Setup of the multi-locus gene-for-gene model

As in classic multi-locus gene-for-gene (GFG) interactions (8; 37; 38), we consider a total of three resistance loci (*L*_1_*, L*_2_*, L*_3_) for both host species and three effector loci (*E*_1_*, E*_2_*, E*_3_) in the pathogen with a strict one-to-one correspondence between host and pathogen loci; alleles from the first resistance locus (*L*_1_) only interact with alleles from the first effector locus (*E*_1_) an so forth. At each resistance locus there are up to two alleles present, namely a susceptible (S) and a resistant (R) allele and each effector locus is characterized by two allelic states, avirulent (A) and virulent (V). Note that here we use the term avirulent (A) to refer to an effector allele that can be recognized by the cognizant resistance allele, but promotes disease in the absence of the corresponding resistance allele (39). For each pair of host and pathogen loci we consider the following GFG-interaction matrix (see Figure 1A):

**Figure 1.**
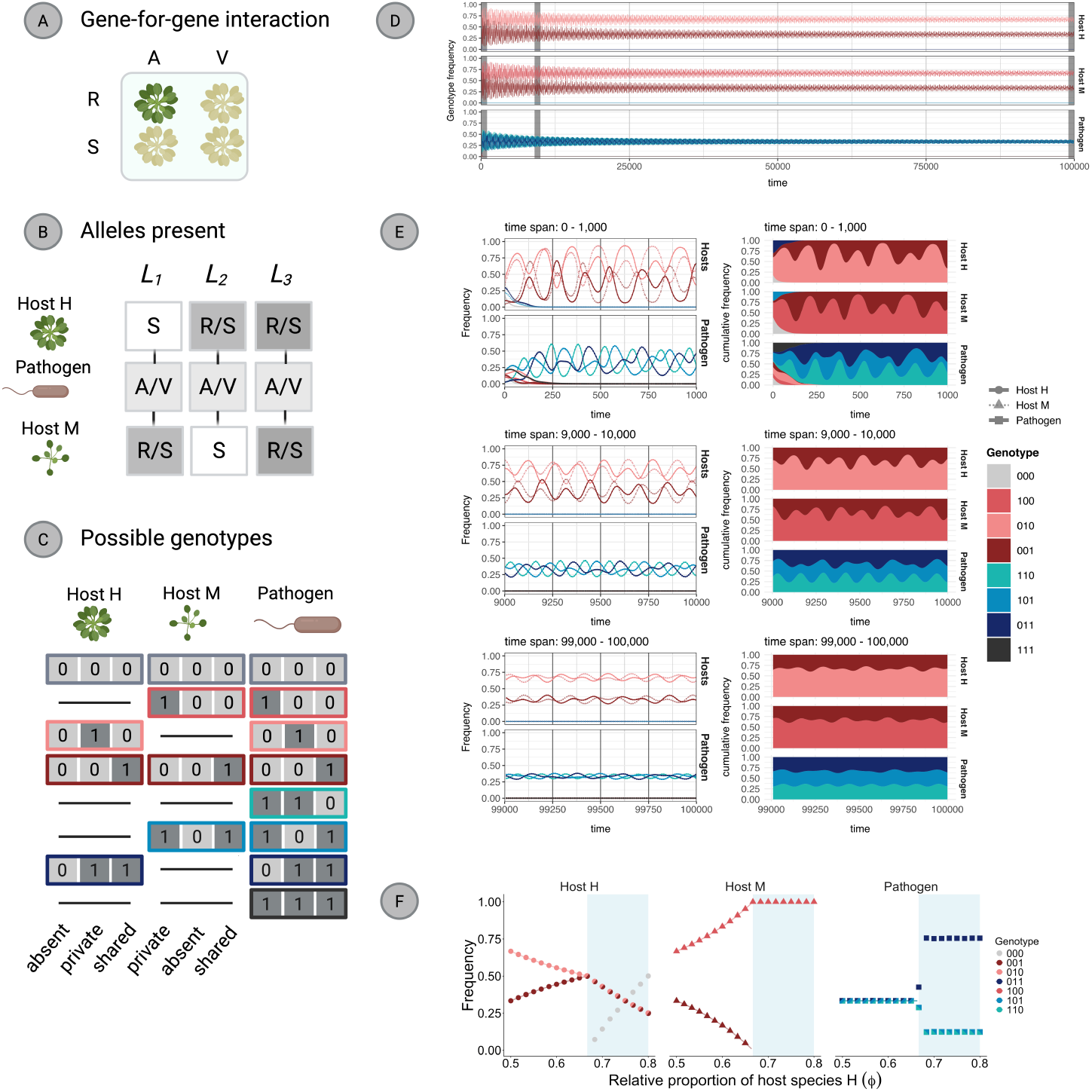
Illustration of the model properties and genotype frequency dynamics for a single simulation (*Ξ_H_* = *Ξ_P_* = 3, *Ω_H_* = *Ω_P_* = 0.1, *ϕ* = 0.5, *β_H_* = *β_P_* = 1, seed = 1, 600, *σ* = 0.85). A: Illustration of the gene-for-gene-model underlying interactions between each pair of host and pathogen loci. Possible host alleles are shown on rows and are R (resistant) and S (susceptible). Pathogen alleles are shown in columns and are A (avirulent) and V (virulent). B: The locus structure of the model. There are a total of three loci in each species, all interacting under the gene-for-gene interaction matrix as shown in A. Host species *H* and *M* share the third locus, while the resistance allele at locus 1 is private to host *M* and the resistance allele at locus 2 is private to host *H*. Possible allelic states in single genotypes are shown in the boxes. As resistance at the first locus is private to host species *M*, all host *H* genotypes are susceptible (S) at this locus. Similarly, as the second locus is private to host species *H*, all host *M* genotypes are susceptible at this locus. C: All possible genotypes in the model for each species. Note that due to the locus structure, certain genotypes cannot be present in both host species. For hosts, 0 corresponds to the susceptible allele at the given locus and 1 reflects the corresponding resistance allele. For pathogens, 0 represents the avirulence allele that can be recognized by a host with a corresponding resistance allele at the locus and 1 depicts a virulence allele at the locus which can escape recognition. D: Full length dynamics of a single simulation for the parameters indicated above. Note that frequencies of pathogen genotypes *P*110*, P*101*, P*011 fluctuate around 1/3 and are hard to distinguish. E: Zoomed in overview of the dynamics for time intervals 0 − 1, 000, 9, 000 − 10, 000 and 99, 000 − 100, 000. On the left, frequencies of all genotypes in both host species are plotted in a single plot (solid lines: host *H*, dashed lines: host *M*). The same dynamics are shown as a cumulative plot for each species on the left. Note that we use the same color codes for host and pathogen genotypes to allow for easier comparisons between the species. F: Frequencies of the different genotypes that coexist in E as a function of the relative proportion of host species *H* (*ϕ*). Lines are predictions from calculating equilibrium frequencies (see Supplementary Information) while shapes are frequencies from simulations averaged over the last 10% of timesteps from a total of 10^5^. Facets and shapes represent different species while colors represent different genotypes. Shapes have multiple colors when multiple genotypes have the same frequency. A-C were created with and all subfigures arranged with BioRender.com.

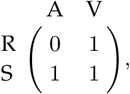

where 1 designates the ability of a pathogen to escape recognition and 0 designates recognition of the pathogen allele (39), which results in decreased growth on the given host. Individual genotypes are denoted as ***x*** = {*x*_1_*, x*_2_*, x*_3_} in host species *H*, ***u*** = {*u*_1_*, u*_2_*, u*_3_} in host species *M* and ***y*** = {*y*_1_*, y*_2_*, y*_3_} in the pathogen.

#### Assigning private and shared resistance to host loci

Our chosen locus structure is based on the observation that *R* genes are often localized in clusters (40; 41; 42) with frequent occurrences of tandem gene duplications (43; 44). To incorporate private and shared resistance between hosts, we assume that resistance locus *L*_3_ is present in both host species (henceforth called the shared locus). Thus, the age of the locus predates the evolutionary split of the two species. Resistance locus *L*_1_ is private to host *M* (hence *x*_1_ = 0 for all host *H* genotypes) and resistance locus *L*_2_ is private to host *H* (hence *u*_2_ = 0 for all host *M* genotypes)(see Figure 1B). Therefore, we assume that independent tandem duplications in each host resulted in the evolution of a new host specific private *R* gene. In summary, this locus structure translates into four possible genotypes in host species *H* (000, 001, 010, 011) and four possible genotypes in host species *M* (000, 100, 001, 101) (see Figure 1C).

#### Fitness expressions

We use differential equations of the following form to describe changes of genotype ***z*** in species *S* (four genotypes in host *H*, four genotypes in host *M* and eight genotypes in the pathogen):

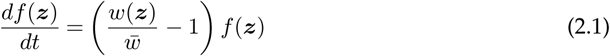

where *f* (***z***) is the frequency of genotype ***z***, *w*(***z***) is its fitness (life-time reproductive success), and *w̄* is the average fitness within the genotype’s species.

The fitness of host ***x*** depends on the probability *Q*(*x, y*) that the host gets infected when interacting with pathogen genotype ***y***, the negative effect of infection on host fitness (*β_H_*), and the baseline fitness *B*(***x***) associated with the genotype. Following (37), we calculate the probability *Q*(***xy***) that host *H* genotype ***x*** gets infected by pathogen genotype ***y*** as a function of the number of effective resistance alleles *d*(***x***, ***y***) (resistance alleles that can recognize the cognizant pathogen allele in a given interaction, see Table S2):

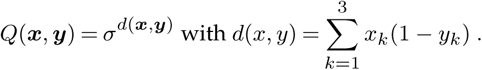

We incorporate the negative fitness effect of infection for the host with a fixed parameter *β_H_* for both hosts and model the positive effect of infection for the pathogen with the parameter *β_P_*. Costs of resistance and virulence are incorporated as a function of the proportion of loci with a resistance (or virulence) allele. The cost functions are characterized by two parameters *Ω* and *Ξ* for each species. *Ω* depicts the maximum loss in fitness when the host (or pathogen) genotype has both resistance alleles (or all three virulence alleles) alleles and is referred to as the maximum cost of resistance (or virulence). The parameter *Ξ* governs the shape of the function that links the number of resistance (virulence) alleles to the corresponding genotypic cost. The baseline fitness (*B*(***z***)) of a genotype (*z*) with *n* resistance (virulence) alleles corresponds to its fitness in the absence of the interacting partner (see Figure S1) and is calculated as follows:

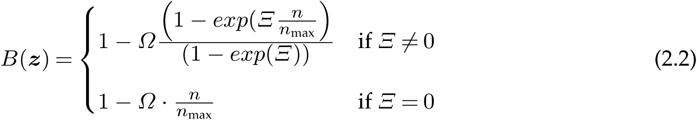

where *n*_max_ is the maximum number possible of resistance alleles in a single host genotype (here *n*_max_ = 2) or the maximum number of virulence alleles possible in a single pathogen genotype (*n*_max_ = 3). If *Ξ <* 0, these costs are decelerating, meaning that the first resistance (or virulence) allele is comparatively expensive, but the addition of subsequent alleles becomes progressively less expensive until the maximum cost is reached. On the other hand, costs are accelerating for *Ξ >* 0, where the first allele is comparatively inexpensive and each additional allele becomes more expensive. Altogether, the fitness expressions can be written as:

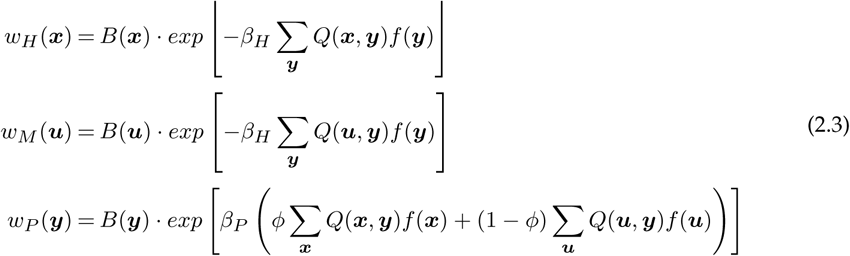

and a detailed overview of variables, parameters and functions can be found in Table 1.

**Table 1:** Description of all parameters and variables in the extended model.

| Name | description |
| --- | --- |
| $K$ | number of interacting pathogen/host loci |
| $x$ | a genotype in host species $H$ |
| $u$ | a genotype in host species $M$ |
| $y$ | pathogen genotype |
| $f(x)$ | frequency of host $H$-genotype $x$ |
| $f(u)$ | frequency of host $M$-genotype $u$ |
| $f(y)$ | frequency of pathogen genotype $y$ |
| $w_H(x)$ | relative fitness of host $H$-genotype $x$ |
| $w_M(u)$ | relative fitness of host $M$-genotype $u$ |
| $w_P(y)$ | relative fitness of pathogen genotype $y$ |
| $d(x, y)$ | number of resistance alleles which recognize an effector in an interaction between host $H$ genotype $x$ and pathogen genotype $y$ |
| $d(u, y)$ | number of recognizing (effective) resistance alleles in an interaction between host $M$ genotype $u$ and pathogen genotype $y$ |
| $\sigma$ | parameter governing how the number of effective resistance alleles relates to the probability of infection |
| $Q(x, y)$ | probability of infection in an interaction between host genotype $x$ and pathogen genotype $y$ |
| $Q(u, y)$ | probability of infection in an interaction between host genotype $u$ and pathogen genotype $y$ |
| $\beta_H = \beta_M$ | effect of infection on host fitness (the same for all host genotypes) |
| $\beta_P$ | effect of a successful infection on pathogen fitness |
| $B(x)$ | relative fitness of host $H$ genotype $x$ in the absence of infection |
| $B(u)$ | relative fitness of host $M$ genotype $u$ in the absence of infection |
| $B(y)$ | relative fitness of pathogen genotype $y$ |
| $\Omega_H = \Omega_M$ | maximum genotypic cost of resistance |
| $\Xi_H = \Xi_M$ | parameter defining the shape of the resistance cost function in host $H$ and host $M$ |
| $\Omega_P$ | maximum genotypic cost of virulence in the pathogen |
| $\Xi_P$ | parameter defining the shape the virulence cost function in the pathogen. |

#### Simulation of the model

We extended the source code from (37) to simulate the dynamics of our system with two host species and the generalist pathogen. We first ran our model assuming equal proportions of both hosts (*ϕ* = 0.5) for 10,000 random combinations of maximum cost of resistance (*Ω_H_*), maximum cost of virulence (*Ω_P_*) and initial genotype frequencies, while fixing the shapes of the cost functions to *Ξ_H_* = *Ξ_P_* = 3 (accelerating costs). Maximum costs of resistance and virulence for individual simulations were independently drawn from a uniform distribution *U* (0.01, 0.3). We randomly initialized the genotype frequencies within a single species by first drawing a random number for each genotype from a uniform distribution U(0, 1) and subsequently normalizing these values so that they sum to 1. Each simulation was run until genotype frequencies did not change appreciably between timesteps or until *T_max_* = 100, 000, whichever occurred first. We considered genotypes to be extinct whenever their frequency dropped below 10^−6^.

For each simulation, we determined (i) the genotypes present at the end of the simulation, (ii) the maximum number of resistance alleles maintained in any host genotype that persisted, (iii) the number and identity of *R*-alleles maintained in each host population, (iv) the polymorphism status of each resistance locus and (v) the genotypes still present in the pathogen population.

Next, we repeated the same procedure for increasing proportions of host species *H*; that is, for *ϕ* ∈ {0.7, 0.9, 1}. Finally, we aggregated results across nine different pairwise combinations of the shapes of the cost functions for the host (*Ξ_H_* ∈ {−3, 0, 3}) and the pathogen (*Ξ_P_* ∈ {−3, 0, 3}) and for four different proportions of host *H* (*ϕ* ∈ {0.5, 0.7, 0.9, 1.0}) (correspondingly, the relative proportion of host *M* is 1 − *ϕ*). Therefore, we jointly analyzed a total of 9 (cost shape combinations) x 4 (relative proportions of host *H*, *ϕ* ∈ {0.5, 0.7, 0.9, 1}) x 10,000 (random combinations of cost maxima *Ω_H_*, *Ω_P_* and initial genotype frequencies) = 360,000 simulations. Throughout all simulations, we fixed the maximum cost of resistance and the shape of the cost function to be the same between the two host species, that is *Ξ_H_* = *Ξ_M_* and *Ω_H_* = *Ω_M_* and report only the values for *Ξ_H_* and *Ω_H_*.

## 3. Results

### (a) Frequency-dependent selection maintains a subset of host and pathogen genotypes with fluctuating frequencies

We start by presenting results for an equal proportion of the two hosts (*ϕ* = 0.5) so that the pathogen encounters both host species with equal probability and for accelerating fitness costs (*Ξ_H_* = *Ξ_P_* = 3). Host-pathogen interactions drive dynamic, reciprocal genotype frequency fluctuations. They also cause the early loss of several genotypes in all three species. (Figure 1E, top).

For the specific choice of maximum genotypic costs (*ω_H_* = *ω_P_* = 0.1), all host genotypes with exactly one resistance allele (shared or private) and all pathogen genotypes with exactly two virulence alleles are maintained (Figure 1E, bottom). Pathogen genotypes with two or three avirulence alleles (*P*_000_*, P*_100_*, P*_010_*, P*_001_) suffer from lower infection efficiency compared to pathogens with a lower number of avirulence alleles, as they can be recognized by at least two host genotypes. The fitness loss due to being recognized outweighs the fitness advantage of having lower genotypic costs and results in the early extinction of strains bearing multiple avirulence alleles (Figure 1E, top). The opposite applies to the pathogen genotype *P*_111_ with three virulence alleles. Although this genotype has the highest overall infection efficiency, it also bears the highest cost of virulence. As host genotypes with two resistance alleles are lost early in the dynamics, pathogen genotypes with two virulence alleles reach almost the same infection efficacy as genotypes with the full complement of virulence alleles (*P*_111_), but at a lower genotypic cost. Completely susceptible host genotypes (with no resistance alleles; *H*_000_ and *M*_000_) rapidly go extinct due to the reduced fitness that they suffer from constant full strength infection.

Once only genotypes that maximize the fitness trade-off remain (around *t* = 700), negative-frequency dependent selection drives fluctuations in their frequencies. Whenever the frequency of the *P*_110_ pathogen increases, the frequencies of *H*_001_ and *M*_001_ (Figure S2a,b) also increase, as these are the only host genotypes able to partially recognize the *P*_110_ genotype. Simultaneously, host genotypes *H*_010_ and *M*_100_ decrease in frequency (Figure S2c,d). As a result of increases in the frequencies of *H*_001_ and *M*_001_, the frequency of *P*_110_ slowly decreases and the frequencies of *P*_011_ and *P*_101_ increase. This, in turn, triggers a decrease in *H*_001_ and *M*_001_ and a corresponding increase in *H*_010_ and *M*_100_ genotypes. Note that these changes are not initially synchronized between the two hosts. Over time, dynamics synchronize and fluctuations dampen (compare Figure 1a,b,c).

### (b) Maximum costs of resistance and virulence determine the number of resistance alleles maintained

Next, we explored the sensitivity of these dynamics to the maximum cost of resistance (*Ω_H_*), the maximum cost of virulence (*Ω_P_*) and the initial genotype frequencies, for fixed shapes of the genotypic cost functions ( *Ξ_H_* = *Ξ_P_* = 3, accelerating costs). As expected, there is a negative relationship between the maximum number of resistance alleles maintained within a single host genotype and the maximum cost of resistance (*Ω_H_* increasing) (Figure S3). However, the maximum number of resistance alleles per genotype depends not only on the maximum cost of resistance but also on the cost of virulence. In particular, the maximum number of resistance alleles maintained within a single genotype generally increases with increasing costs of virulence. These results highlight previous findings that host genotypic costs and pathogen genotypic costs (8; 37; 38) jointly determine the number of resistance alleles that persist in the repertoire of a single host genotype.

### (c) Host proportion alters the maintenance of host and pathogen genotypes

Returning to the dynamics in Figure 1, we explore how the relative proportion of host species *H* modifies which genotypes coexist while fixing the maximum costs of resistance and virulence. Analytical formulas for the equilibrium frequencies of each host and pathogen genotype (lines in Figure 1F) predict the averages of numerical simulations (points in Figure 1F) well. Our equilibrium analysis of this particular set of maximum costs reveals that increasing the relative proportion of host *H* increases the frequency of the shared resistance genotype in host *H* and decreases the frequency of the private resistance genotype (Figure 1F). Conversely, a larger proportion of host species *H* favors private or shared resistance in the less common host species *M* (Figure 1F). Our theory predicts that the private resistance genotype in host species *M* is excluded at *ϕ* = 2*/*3, and numerical simulations confirm this prediction. Interestingly, when *ϕ >* 2*/*3, the genotype of host species *H* with no resistance alleles can invade and the pathogen frequencies shift to fluctuate around a new equilibrium state. Therefore, modifying only the relative proportion of the two host species can induce changes in which genotypes are maintained across all three species.

### (d) The less abundant host preferentially maintains the resistance allele at the private locus

Taking a simulation approach, we next assessed how the relative proportion (*ϕ*) of host species *H* interacts with the costs of resistance (*Ω_H_*) and virulence (*Ω_P_*) to determine the dynamics at resistance loci. We first ask which resistance alleles are maintained and then determine whether the resistance alleles fix in the population or persist as a polymorphism with susceptible alleles. In our haploid model, the latter implies that there are at least two genotypes maintained in the population that differ in their allelic state at a given locus. We begin by presenting results on the maintenance of resistance alleles and consider the maintenance of polymorphism in the next subsection.

When both hosts are equally abundant, private and shared resistance alleles are maintained for a wide range of maximum cost combinations (Figure 2a and b) in both host species. The propensity to maintain resistance alleles persists in the more abundant host *H* (increasing *ϕ*) as its relative frequency increases (Figure 2a,b right, top to bottom), although there is an expansion in the proportion of parameter space (combinations of maximum genotypic costs *Ω_H_* and *Ω_P_*) in which both resistance alleles are lost. In contrast to the more abundant host *H*, the less abundant host species *M* tends to retain only the private resistance allele and lose the shared resistance allele (Figure 2a,b left, top to bottom). These patterns are considerably robust across shape functions (Figure 2c). Importantly, they are also consistent with our analytical predictions from Figure 1F. An additional interesting observation is that there are only a few cases where the shared resistance allele persists alone, in the absence of the private resistance alleles (light blue range in Figure 2c). When the pathogen interacts exclusively with host species *H* (*ϕ* = 1), we observe equal proportions of simulations in which only the private or only the shared resistance allele is maintained because, in the absence of host *M*, the shared and the private resistance loci in host *H* become equivalent. We note that for some part of the parameter space, there is an effect of the initial genotype frequencies on the final outcome of the interaction (for one specific case see Figure S6). Simulations with fixed and equal initial frequencies illustrate a complex landscape of possible equilibrium states (Figure S8) as a function of the costs of resistance and virulence.

**Figure 2.**
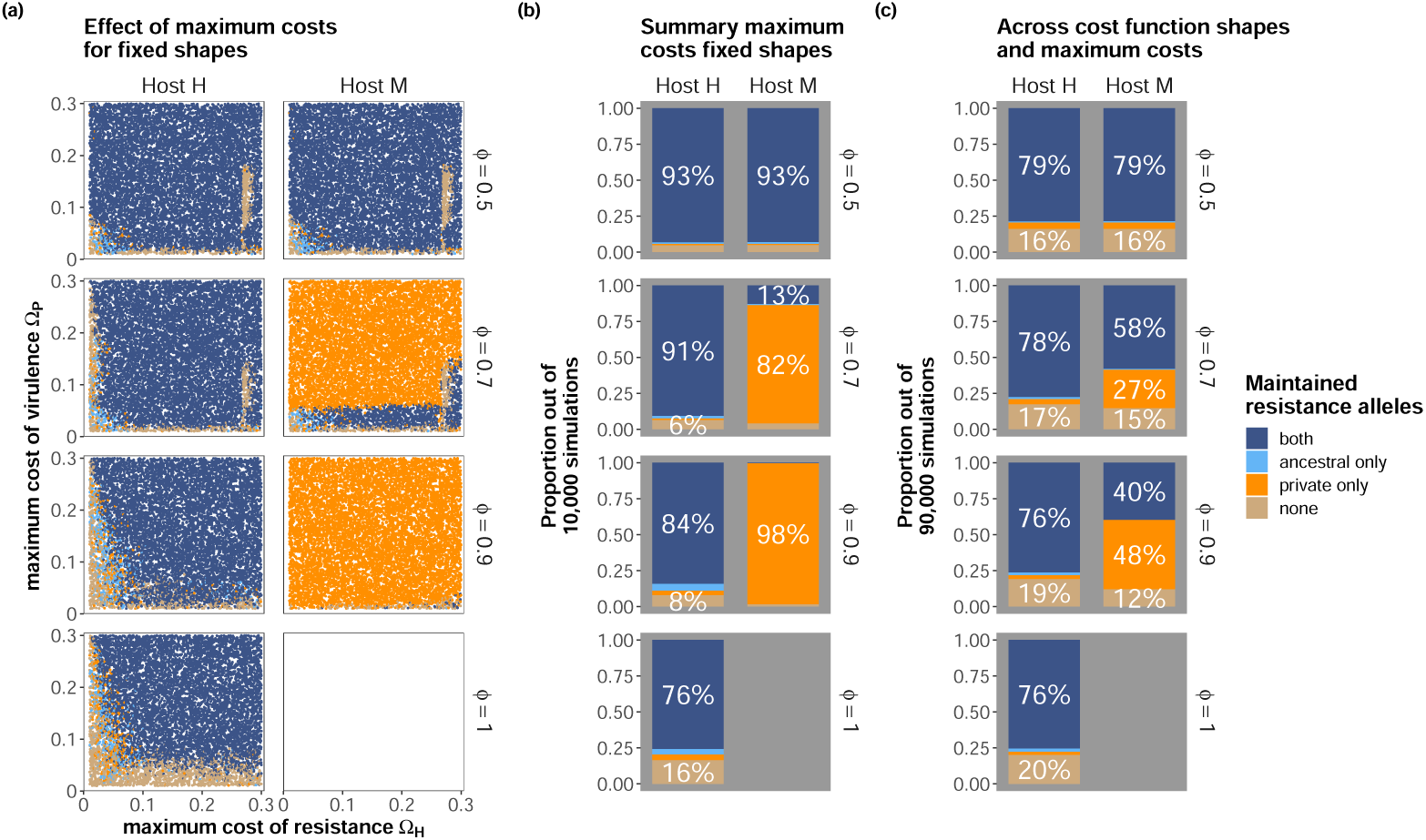
Overview on resistance alleles that are maintained in both host species. (a) and (b) Results for fixed shapes of the cost functions (*Ξ_H_* = *Ξ_P_* = 3) and 10,000 simulations with random initialization of all genotype frequencies and the value for the maximum cost of resistance and virulence drawn from *Ω_H_* = *Ω_M_* ∼ U(0.01, 0.3) and *Ω_P_* ∼ U(0.01, 0.3). (c) Results across all pairwise combinations of*Ξ_H_* = *Ξ_M_* ∈ {−3, 0, 3} and *Ξ_P_* ∈ {−3, 0, 3} with random initialization of all genotype frequencies and the value for the maximum cost of resistance and virulence drawn from *Ω_H_* = *Ω_M_* ∼ U(0.01, 0.3) and *Ω_P_* ∼ U(0.01, 0.3). (a) Shows the detailed results for the fixed shapes for random combination of *Ω_H_* (x-axis) and *Ω_P_*. These results are summarized as stacked barplots in (b). Note that only relative proportions ≥5% are labelled. The relative proportion of host species *H* increases from top to bottom. Panels (c) shows the summarized results across all nine pairwise combinations of the shapes of the cost functions and randomly drawn combinations of *Ω_H_*, *Ω_P_* and initial genotype frequencies as outlined above. Results are shown for the same set of simulations as in Figure 3.

### (e) Fixation of the private resistance allele is favored in the less common host

From an evolutionary perspective, both the persistence of a resistance allele and the maintenance of a polymorphism at that allele are of interest. When hosts are equally frequent (*ϕ* = 0.5), polymorphism is observed for both private and shared resistance loci for a wide range of maximum cost combinations (see Figure 3a and b top left). Polymorphisms in the private loci (*L*_1_ for host *M*, *L*_2_ for host *H*) are lost when the cost of resistance is low and the cost of virulence is intermediate to high (see Figure 3a), because under these conditions the host fixes for the resistance allele at the private locus (Figure S4a, top). However, the polymorphism at the shared locus persists in both hosts under these same conditions.

**Figure 3.**
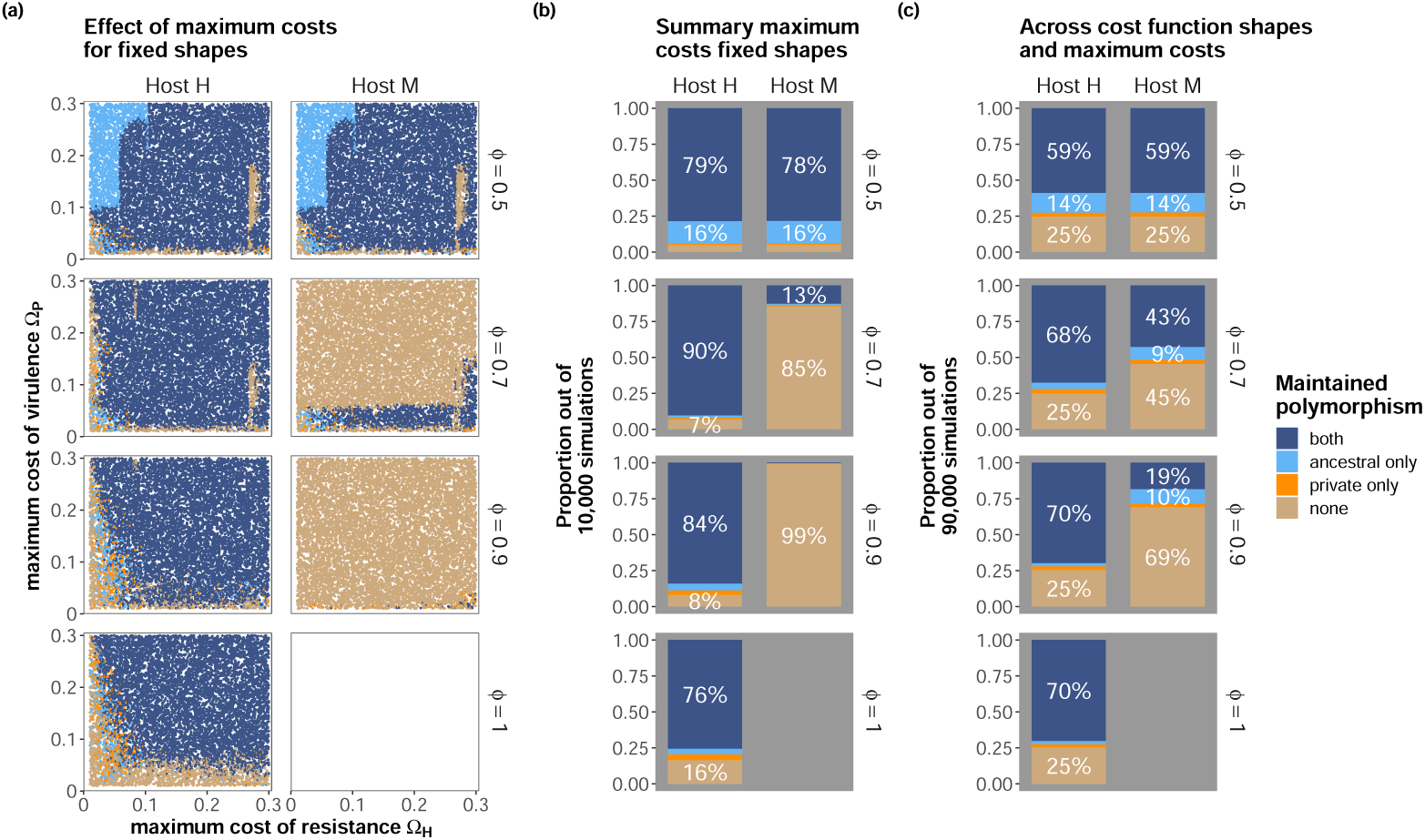
Overview on polymorphism that is maintained at resistance loci. (a) and (b) Results for fixed shapes of the cost functions (*Ξ_H_* = *Ξ_P_* = 3) and 10,000 simulations with random initialization of all genotype frequencies and the values of the maximum cost of resistance and virulence drawn from *Ω_H_* = *Ω_M_* ∼ U(0.01, 0.3) and *Ω_P_* ∼ U(0.01, 0.3). (c) Results across all pairwise combinations of*Ξ_H_* = *Ξ_M_* ∈ {−3, 0, 3} and *Ξ_P_* ∈ {−3, 0, 3} with random initialization of all genotype frequencies and the value for the maximum cost of resistance and virulence drawn from *Ω_H_* = *Ω_M_* ∼ U(0.01, 0.3) and *Ω_P_* ∼ U(0.01, 0.3). (a) Shows the detailed results for the fixed shapes for random combination of *Ω_H_* (x-axis) and *Ω_P_*. These results are summarized as stacked barplots in (b). Note that only relative proportions ≥5% are labelled. The relative proportion of host species *H* increases from top to bottom. Panels (c) shows the results summarized across all nine pairwise combinations of the shapes of the cost functions and randomly drawn combinations of *Ω_H_*, *Ω_P_* and initial genotype frequencies as outlined above. Results are shown for the same set of simulations as in Figure 2.

As the proportion of the two hosts becomes more skewed (Figure 3 top to bottom), we find that the less common host *M* tends to lose polymorphism at both the private and shared resistance loci (Figure 3 left columns in subfigures) because the private locus becomes fixed for the resistant allele and the shared resistance allele is lost (see Figure 2, S4 left column, top to bottom). In contrast to the minor host, the more common host species *H* maintains polymorphism for both resistance loci for a wide range of parameter combinations (Figure 3ab, S4ab right columns in subfigures). These overall patterns hold for a wide range of maximum costs and shapes of the cost functions (Figure 3c, S4c), and are consistent with our analytical predictions for the scenario in Figure 1.

### (f) Polymorphism is more likely to be maintained at the shared relative to private loci, promoting trans-specific polymorphism

Across a range of combinations of the maximum costs of resistance and virulence, and shapes of the cost functions, we find frequent instances of polymorphism at the shared resistance locus, especially for the more common host species (Figure 3b, S4, S5). In particular, we find that for the more abundant host species *H*, the most frequent outcome is the maintenance of polymorphism at both resistance genes (private and shared) (Figure 3b). Remarkably, trans-specific polymorphism at the shared resistance gene tends to be maintained, even with an increasing imbalance in the frequencies of the two host species (increasing *ϕ*) (Figure S7). Polymorphism exclusive to private resistance genes is much rarer than polymorphism that is exclusive to the shared resistance locus.

### (g) The pathogen adapts to the more common host species

The fitness of pathogen genotypes is affected by the relative proportion of both host species (through the parameter *ϕ*). We therefore asked how co-evolutionary dynamics change as the relative abundance (*ϕ*) of host *H* increases. A frequency increase of host *H* (*ϕ >* 0.5) can be interpreted as either: (i) host *H* has a higher abundance compared to host *M*, or (ii) there is some direct or indirect (i.e. through a transmitting vector) preference of the pathogen for host *H*. We were specifically interested in whether we would observe a shift in the pathogen genotypes that are maintained as a result of more frequent interactions with host *H*. Averaging over different combinations of the maximum costs *Ω_H_* and *Ω_P_* and different shapes of the cost functions, we find a decrease in the probability that a pathogen genotype with full virulence (*P*_111_) persists (blue colors) as interactions with host *H* become more frequent (Figure 4 right; from top to bottom). The reduced probability of maintaining the fully virulent pathogen is accompanied by an increase in the probability that genotypes with two virulence alleles are maintained (pink/red colors).

**Figure 4.**
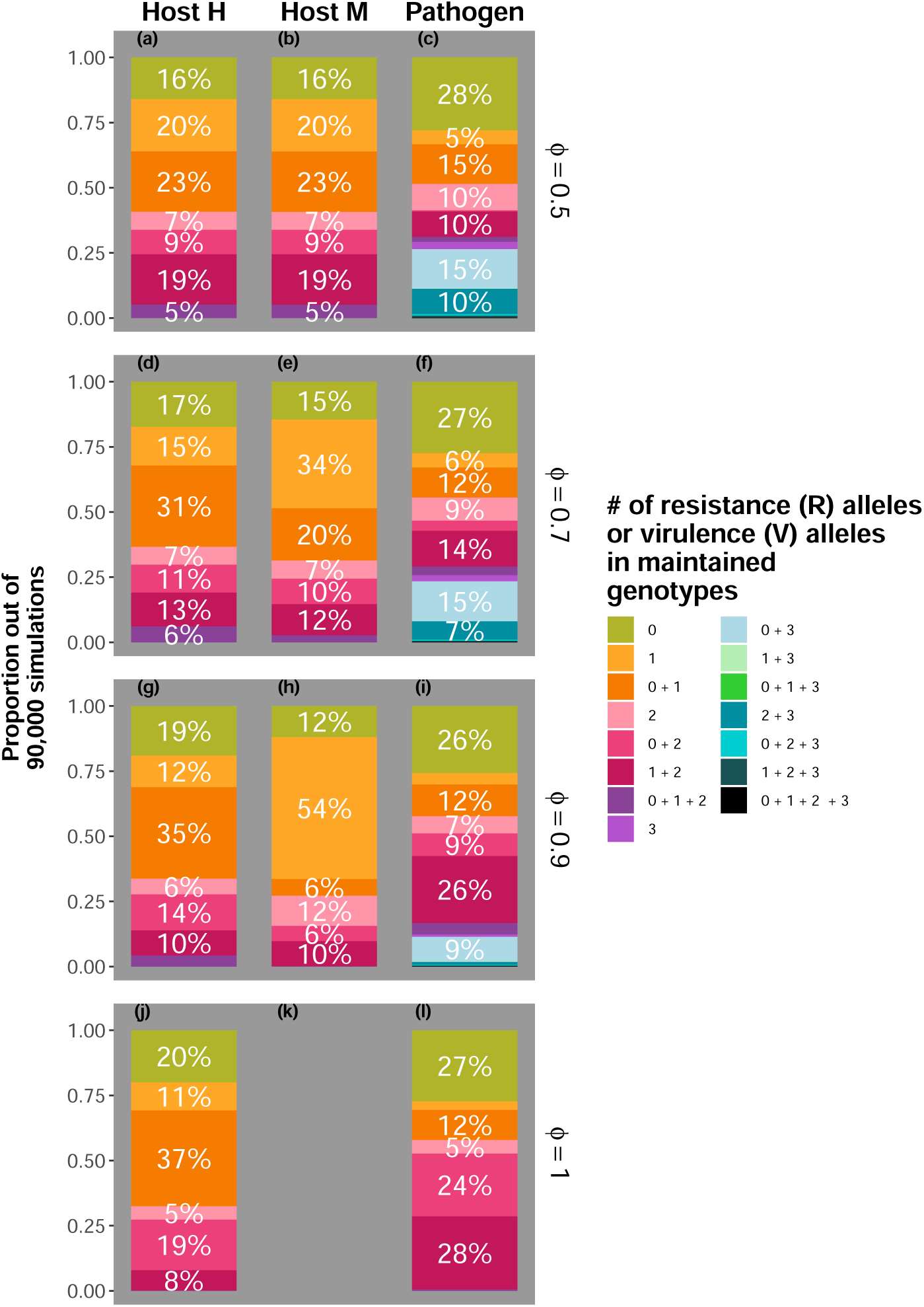
Genotype combinations that are maintained in host *H*, host *M* and the pathogen *P* (columns) for a wide range of cost combinations and shapes depending on the relative proportion *ϕ* (rows) of host *H*. The legends are to be read as follows: A single number means that only genotypes with the corresponding number of resistance **R**/virulence **V**) alleles have been maintained in the given species; several numbers combined with a ‘+’ indicates that genotypes with different numbers of resistance *R*/virulence *V* alleles are still present at the end of the simulation. Note that the color codes in the hosts and the pathogen are the same for combinations that can occur in hosts and pathogens. Also note that there might be several genotypes with a given number of R-alleles/V-alleles maintained. Each stacked bar corresponds to a total of 90,000 simulations with 10,000 simulations each for all pairwise combinations of *Ξ_H_* = *Ξ_M_* ∈ {−3, 0, 3} and *Ξ_P_* ∈ {−3, 0, 3}. The 10,000 simulations for each combination of *Ξ_H_* = *Ξ_M_*, *Ξ_P_* and *ϕ* were run by randomly drawing the initial frequencies of all genotypes, *Ω_H_* = *Ω_P_* ∼ U(0.01, 0.3) and *Ω_P_* ∼ U(0.01, 0.3). All simulations where summarized at *t* = 100, 000.

The alternative hosts, *H* and *M* reveal some contrasting patterns. As the frequency of host *H* increases (Figure 4 left top to bottom), we see an expansion in the parameter space for which host species *H* carries zero (*H*_000_) and one (*H*_010_ and/or *H*_001_) resistance allele and an expansion in the parameter space for which host *M* carries exactly one resistance allele (Figure 4 middle, top to bottom). In many instances, retention of genotypes with only one resistance allele in host *M* corresponds to fixation of the private resistance allele (*M*_100_) (Figure 1F, 2, S4, S5). Taken together, our results suggest that as interactions with host *H* become more frequent, the pathogen tracks genotype frequencies in this host more closely with the consequence that selection favors genotypes that maximize fitness on host species *H*.

## 4. Discussion

Understanding the proximate causes that maintain high levels of diversity and polymorphism at plant resistance genes has been a longstanding research problem. Recent evidence suggests that the broader community context in which these interactions occur may play an important role (14; 33; 45). In line with these results, we show that interactions of closely related hosts with a shared generalist pathogen contribute to the maintenance of resistance alleles and resistance gene polymorphisms. Our results further highlight that evolutionary dynamics at shared and private resistance genes are contingent on the relative abundances of the host species, along with the genotypic costs of both resistance and virulence. Therefore, our model is consistent with previous models demonstrating that species-specific trade-offs play a prominent role in shaping reciprocal evolutionary dynamics. (8; 9; 13; 37; 38; 46). In addition, our results suggest that trans-specific *R* gene polymorphisms may be promoted by interactions with a shared generalist pathogen, rather than being contingent on any particular functional attribute of the respective *R* gene. However, the extent may depend on the specific cost functions and relative abundances of the two host species.

### The evolution of *R* gene repertoires depends on host-specific and pathogen specific fitness trade-offs

In line with previous findings (8; 9; 13; 37; 38; 46), we show that the evolutionary dynamics of *R* genes are determined not only by the cost of resistance, but also by the cost of virulence in the pathogen. While these results may seem counterintuitive at first, they are a direct result of the frequency of pathogen alleles driving the selective pressures on host genotypes and vice-versa (termed indirect negative-frequency dependent selection in the literature (9)). If virulence is costly for the pathogen, and hence the frequency of avirulent genotypes increase, resistance in the host will be favored, provided that the advantage of resistance sufficiently outweighs its associated costs. Thus, our results underscore the importance of an integrative consideration of host-specific and pathogen-specific trade-offs for understanding the evolutionary dynamics of resistance genes and repertoires.

### A better understanding of cost functions in both hosts and pathogens is required for a general understanding of pathogen-host dynamics

We observed variable evolutionary outcomes depending on the specific costs of resistance and virulence, highlighting the importance of these costs for discerning the evolution of resistance and effector gene repertoires. Measuring gene-specific costs of resistance and virulence has proven to be challenging, due to their dependence on the ecological and/or genomic context (10; 13). While studies of single genes have offered useful insights into the magnitude of costs and the extent to which they can be alleviated through pleiotropic effects and genomic context (14; 15), our results underscore the importance of determining the costs associated with entire repertoires for understanding the evolution of full complements of resistance genes or effectors in host and pathogen genomes.

A recent theoretical study (47) suggests that accelerating cost-benefit trade-offs may be a valid first assumption. Empirically, Huang et al. (48) found evidence for both decelerating and accelerating growth-defense trade-offs in a bacterial prey, depending on whether the prey interacted with a co-evolved ciliate or an ancestral ciliate predator. In a study of *P. infestans* isolates, Montarry et al (2010) (49) found a negative correlation, best characterized as linear or accelerating, between the number of virulence factors and overall fitness, when infecting susceptible potato cultivars. Sequentially mutating functional effectors in the *Xanthomonas axonopodis* pv. *vesicatoria* strain Wichmann and Bergelson (12) revealed complex, but sometimes additive effects, and a dominant role of one particular effector. Overall, these examples demonstrate that cost functions vary across systems, which motivates our exploration of various cost functions.

### Pathogen maladaptation to a less frequent host is a result of the pathogen adapting to the more common host species

One commonly held prediction in the local adaptation literature (35; 36) is that pathogens tend to be ahead of the host in co-evolutionary interactions due to their shorter generation times and higher mutation rates. By investigating patterns of resistance of *A. thaliana* accessions to resident and non-resident strains of the generalist pathogen *P. syringae*, Kniskern et al. (34) found significantly higher levels of resistance to local versus non-resident pathogen strains. They suggested that this may be a consequence of pathogens adapting to more commonly encountered hosts, such as crops grown nearby, thereby resulting in pathogen maladaptation to the less frequently encountered host. Our results provide some evidence for this hypothesis. First, we observe a tendency in the less common host to fix for the private resistance allele. Second, we see an overall shift in the pathogen towards maintaining genotypes with a lower number of virulence alleles. In many cases, this corresponds to a loss of the virulence allele that escapes recognition by the private resistance allele in the less common host. Thus, interactions of a generalist pathogens with a less dominant host such as *A. thaliana*, provide an additional mechanism to promote pathogen maladaptation (for alternative explanations, see 36; 50; 51).

### The community context matters for the evolution of resistance genes

Analyses of *R* gene polymorphism patterns have revealed diverse evolutionary histories (41; 52). In many cases, these histories are more consistent with trench-warfare dynamics (17), which are characterized by sustained allele frequency fluctuations of several alleles, rather than arms-race dynamics (16), in which alleles are recurrently fixed. Theoretical studies of simple host-pathogen interaction models (one host species tightly co-evolving with one pathogen species) have shown that polymorphism often tends to be transient (reviewed in 9), but can be promoted by direct negative frequency-dependent selection (dNFDS), whereby the fitness of an allele/genotype depends on its own frequency. Mechanisms that can generate dNFDS include epidemiological dynamics and asynchrony between host and pathogen life-cycles (3; 9). Unable to explain the ancestral presence/absence polymorphism of the *A. thaliana* resistance gene *Rps5* as a result of tight co-evolutionary interactions with *P. syringae*, Karasov et al. (14) proposed that diffuse host-pathogen interactions may maintain this *R* gene polymorphism. Here, we have found further indication that community interactions, especially those between closely related hosts and generalist pathogens, may play a role in maintaining polymorphism by creating an imbalance in the intensity of selection. This is due to the fact that the pathogen species is confronted with two hosts whereas each host species interacts with only one pathogen. Our results thus reaffirm that it is advantageous to shift our focus from tightly coupled interactions to community-wide context.

### Current limitations of the model

While our model revealed an interesting mechanism that promotes the maintenance of polymorphism, several directions remain for future research. First, our model assumes haploid host and pathogen species. The assumption of haploid genotypes accurately describes bacterial pathogens, however many relevant plant pathogens (53) are oomycetes that are characterized by complex life-cycles. When considering the host, we consider our assumption of haploid hosts to be valid for *Arabidopsis*, which is a diploid plant with such a high selfing rate that most loci are effectively homozygous. Nevertheless, not all plants would behave as haploids. Indeed, outcrossing and higher ploidy levels are common in plants (54). Mating systems and ploidy levels may affect dynamics due to the importance of dominance and the interaction between genotypic costs and ploidy levels. Heterozygote advantage may play a further powerful role in maintaining diversity. Further, recombination in out-crossing species can reintroduce extinct genotypes, likely maintaining polymorphism more extensively, with specific outcomes depending on the difference between host and pathogen recombination rates. Similarly, hosts may have different generation times. As we study a continuous-time model we argue that unless these differences lead to a separation of time-scales they should not qualitatively alter the results.

Currently, our model is based on a very short repertoire that includes a maximum of two functional resistance genes per host and three effectors in the pathogen. Future work should explore the effect of larger repertoire sizes, but as emphasized earlier, this modelling extension will require a better understanding of how genotypic costs link to repertoire size. Furthermore, effector recognition by plant resistance genes may take various forms and is better described by a network than a set of strict one-to-one interactions (55; 56; 57; 58). It will be interesting to understand how such a network structure alters co-evolutionary outcomes. Models that better capture the immune network will ultimately be necessary to advance our understanding of how resistance gene repertoires evolve.

We consider a very specific architecture of the resistance and effector genes, and do not account for reintroduction of previously extinct genotypes as a result of recombination. Therefore, our model best reflects a resistance gene cluster where each host species has separately gained a new resistance gene through a process such as tandem gene duplication and a pathogenicity island in the pathogen (59; 60). However, *R* gene clusters in plants are known to be highly dynamic (44) and subject to both recombination and gene conversion (43; 61). The importance of recombination on resistance gene evolution has been shown in a model (45) demonstrating that the evolution of broad-spectrum and specific-resistance genes is contingent on the amount of recombination.

Finally, our model is based purely on genotype frequencies and neither incorporates the effect of genetic drift nor eco-evolutionary feedback (host-pathogen, host-host). Hence, we implicitly assume that the population sizes of the hosts and the pathogen are time-invariant and large. Eco-evolutionary feedback can create direct frequency dependent selection, which generally promotes stable maintenance of polymorphism (2; 3; 9; 62; 63). Thus, we speculate that accounting for eco-evolutionary feedbacks is unlikely to result in reduced amounts of genetic polymorphism. On the other hand, genetic drift, (that is, random changes in allele frequencies), can alter allele frequency trajectories, sometimes even causing the loss of a deterministically stable polymorphism (64; 65). Nonetheless, our simulations reveal consistent invasion of coexisting genotypes from low frequencies, even though the equilibria we identify are only neutrally stable, suggesting that, at least in this model, genotypes may be partially buffered against drifting to exclusion. However, this may be no longer true when host species experience very strong and recurrent population bottlenecks. Investigating the effect of higher host and pathogen ploidy levels, recombination, extended *R* gene repertoire-effector repertoire interactions, alternative life-cycles and demographic changes in the host species, are thus important extensions for future work.

### Applicability to other host-pathogen systems

While we focused on interactions between plant hosts and pathogens in our model, our results have interesting implications for other host-pathogen systems. First, the evolutionary dynamics and the maintenance of polymorphism at genes involved in immune responses may be contingent on how frequently pathogens encounter alternative hosts. Second, the observed trans-specific polymorphism at immune genes in hosts such as among *Daphnia* species and between humans and apes (26; 29; 30) may have similarly arisen from interactions with generalist pathogens. That said, determining the extent to which our results apply to other systems will require future work. For example the genetic architecture underlying the interaction between *Daphnia* and pathogens may be better described by a matching-alleles model (24). Thus, future models will need to account for such differences in recognition architecture. As pointed out in the previous section, additional assumptions regarding ploidy and recombination may warrant future investigation to understand their specific implications. Thus, we hope that our model encourages the investigation of systems that go beyond plant host-pathogen interaction models.

## 5. Conclusion

Our results show how interactions between a shared generalist pathogen and two related host species can shape their resistance gene evolution and the extent to which resistance alleles and *R* gene polymorphism are maintained. This offers a new perspective for our understanding of the evolution of resistance gene repertoires and allelic diversity, and how shared trans-specific polymorphism may be maintained as a by-product of such interactions, rather than any particular functional attribute of the respective *R* gene. As such, the results of our model potentially help to explain the maintenance of trans-specific polymorphism at immune genes in various systems (26; 28; 29; 30).

While our results are quite robust across genotypic functions, specific outcomes depend on the maximum costs and the shape of the cost functions. This underlines the need for obtaining a more refined understanding of how single members of resistance or effector gene repertoires contribute to the overall fitness of genotypes. We further show that as interactions of the pathogen with a host species become less frequent, pathogen adaptation to genotypes in the more common host species frees the less common host species to fix for private resistance alleles. Overall, our work provides a theoretical understanding of multi-host-pathogen interactions and highlights that community context is a crucial factor shaping the evolution of host-pathogen interactions.

## Acknowledgements.

The authors thank Mercedes Pascual and Enrique Rojas for helpful discussions and Aurélien Tellier and McCall Calvert for useful comments on earlier versions on the manuscript. This work was supported by Tamkeen under the NYU Abu Dhabi Research Institute Award to the NYUAD Center for Genomics and Systems Biology (ADHPG-CGSB) to JB, and grant SFI-PD-Grant-01308072 of the Simons Foundation to JB. TLG was supported by Schmidt Science Fellows, in partnership with the Rhodes Trust.

## Data availability

All codes for simulating the model and data underlying the figures are available at https://doi.org/10.5281/zenodo.15270443 and https://github.com/ PopRGen/TSPolyR2.git.

## A. Supplementary information

### (a) Stability of equilibria

When considering the dynamics of our model, we can eliminate one genotype from each host and the pathogen frequencies because they must sum to one. The Jacobian evaluated at any equilibrium has the form

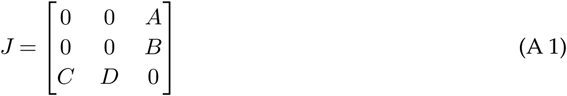

where 0 denotes a matrix of 0s of the appropriate size, *A* encodes how the change in host *H*’s frequencies depend on the pathogen genotypes and *B* is the same quantity but for host *M*. *C* and *D* describe how pathogen frequencies depend on host *H* and *M* respectively. Because this matrix has a block structure with only zeroes in the diagonal block, its eigenvalues come in pairs reflected over the real axis i.e. if *λ* is an eigenvalue of *J* then so is −*λ*. As a result, no equilibrium of this model is stable, although equilibria can be neutrally stable in which the eigenvalues have zero real part. Neutral stability likely leads to the persistent fluctuations we observe in Figure 1.

### (b) Deriving the symmetric polymorphism

Let us consider the kind of polymorphism we observed in Figure 1. We will show that at equilibrium, maintained genotypes within species have the same fitness. We perform this calculation for the set of host and pathogen ranges that we know to be maintained for the chosen costs (Figure 1). In this case, the relevant genotypes are *x* = 010, 001, *u* = 001, 100 and *y* = 011, 101, 110. Because each host (or pathogen) has the same number of resistance (or virulence) alleles, their genotypic costs are equivalent. For host *H*, we have

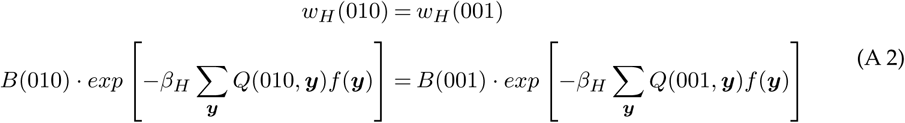

which is true when

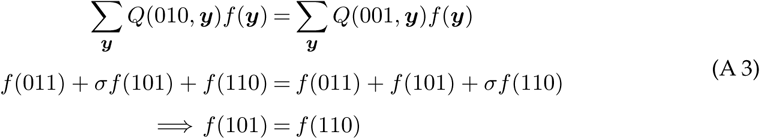

so we know those two pathogen equilibrium frequencies are equal. The same analysis for host *M* shows that *f* (110) = *f* (011) which, when combined with the sum constraint that *f* (011) + *f* (101) + *f* (110) = 1, proves that *f* (011) = *f* (101) = *f* (110) = 1*/*3. In other words, whenever this state is stable to invasion by other host and pathogen ranges, the pathogen frequencies are all equal. Now, we analyze pathogen fitness. Following the same logic as in the host case and carrying out the algebra, we have that

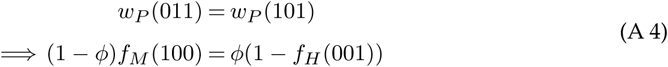

While

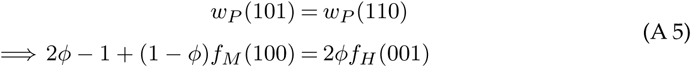

so that

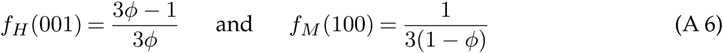

at equilibrium. For *ϕ* ≥ 2*/*3, *f_M_* (100) ≥ 1 so *f_M_* (001) ≤ 0 and the polymorphism in host *M* does not occur as we observe in Figure 1F.

## B. Supplementary Figures

**Figure S1.**
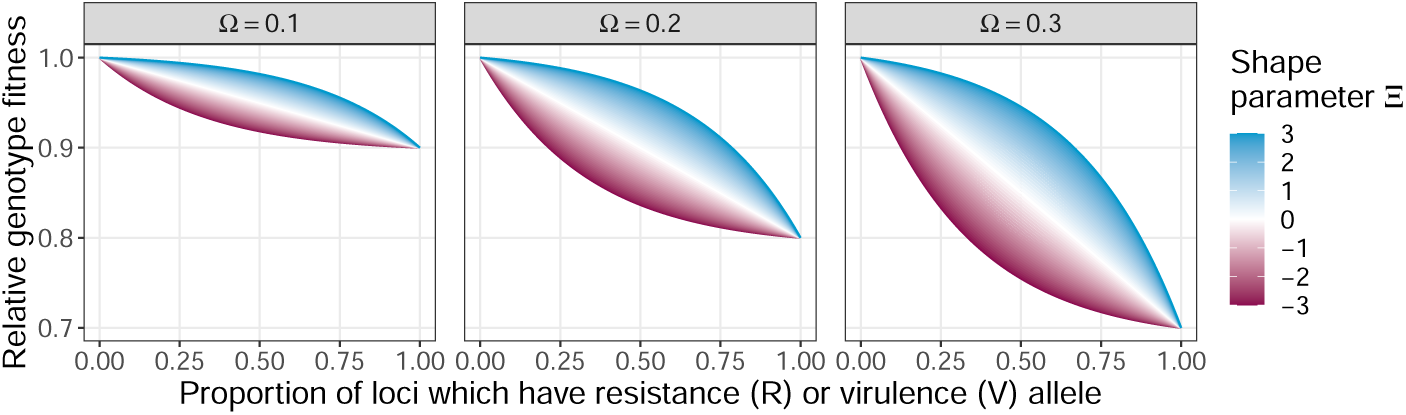
Fitness costs. Relationship between the proportion of loci with a resistance (R-allele)/virulence (V-allele) (x-axis) and fitness for different combinations of *Ω* and *Ξ*. Here *Ω* is the maximum reduction in fitness when a given genotype in species *S* has all possible resistance or virulence alleles and *Ξ* governs the shape of the function.

**Figure S2.**
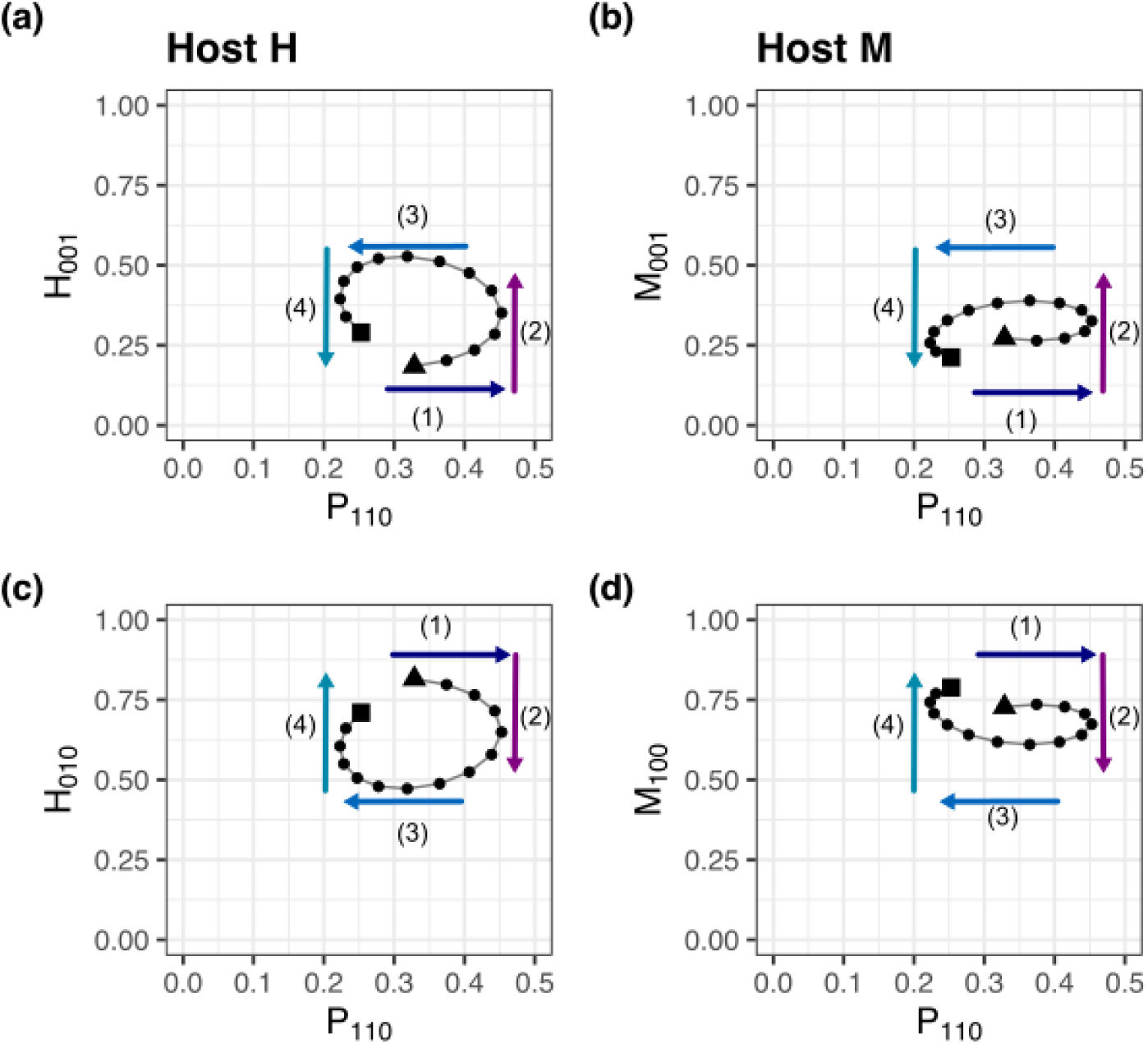
Phase plane plots for the changes of different host genotypes (y-axis) and the pathogen genotype *P*110 (x-axis) for the time interval *t* = 9120 to 9150 for the dynamics shown in Figure 1D,E. Consecutive time points are connected by segments. The first time point in the interval is marked by a triangle, the last one by a square. Results are shown for: *Ξ_H_* = *Ξ_M_* = *Ξ_P_* = 3, *Ω_H_* = *Ω_M_* = *Ω_P_* = 0.1, *ϕ* = 0.5, *β_H_* = *β_P_* = 1, seed = 1, 600, *σ* = 0.85.

**Figure S3.**
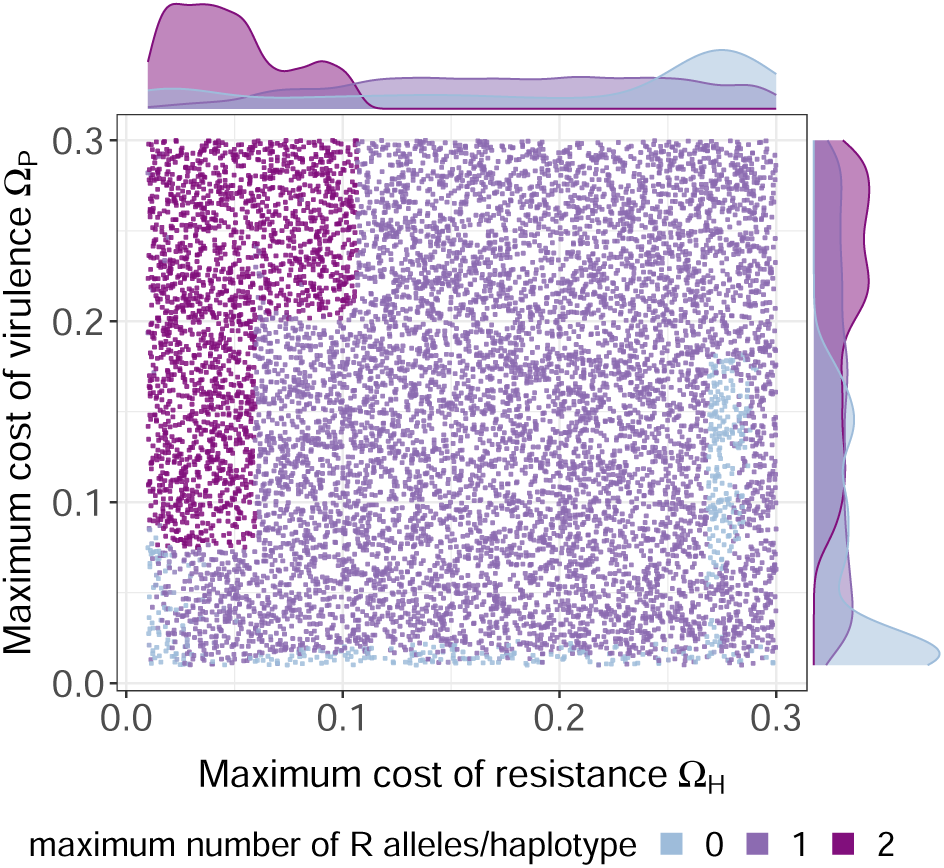
Maximum number of resistance alleles maintained in a single genotype of host species *H* for different combinations of the maximum costs of resistance (*Ω_H_*) and virulence (*Ω_P_*) based on 10,000 randomly drawn combinations of *Ω_H_* ∈ U(0.01, 0.3) and *Ω_P_* ∈ U(0.01, 0.3) and initial genotype frequencies. Note we here show the genotype with the highest number of resistance alleles maintained in host *H* (that is 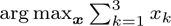). This does not exclude the possibility that additional genotypes with fewer resistance alleles have been maintained. The marginal probability densities for maintaining different maximum numbers of resistance alleles (that is 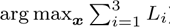) per genotype for increasing costs of resistance *Ω_H_* are shown and for different maximum values of virulence *Ω_P_* on the left. Each point in the main plot shows the maximum number of resistance alleles maintained in a single host genotype for a random combination of *Ω_H_* (x-axis) and *Ω_P_* (y-axis). Points are colored according to the maximum number of resistance alleles found in any maintained host genotype (i.e if genotypes *H*010 and *H*011 are maintained the point would be colored in dark purple).

**Figure S4.**
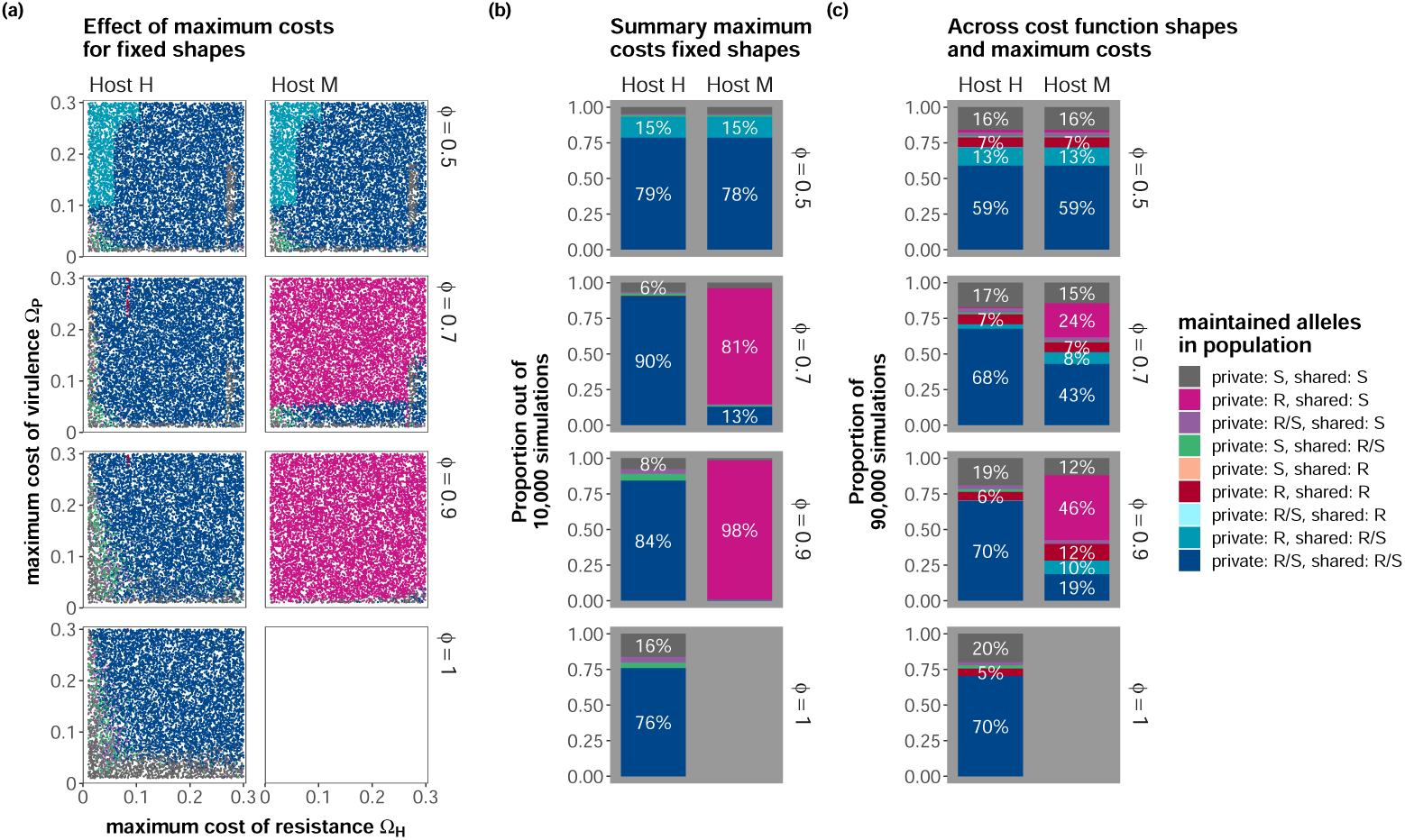
Type of alleles maintained in host species *H* and *M* with the relative frequency of host species H (*ϕ*) increasing from top to bottom in each subfigure. (a) For different combinations of the maximum costs of resistance (*Ω_H_* =*Ω_M_*) and virulence (*Ω_P_*) based on 10,000 randomly drawn combinations of *Ω_H_* = *Ω_M_* ∈ U(0.01, 0.3) and *Ω_P_* ∈ U(0.01, 0.3) and initial genotype frequencies for *Ξ_H_* = *Ξ_P_* = 3 for fixed shapes of the genotypic cost functions (*Ξ_H_* =*Ξ_M_* =*Ξ_P_*). (b) A summary of the results shown in (a) (only frequencies *>* 5% are labelled). (c) Results averaged over nine different combinations of the shape parameters. Results are shown for the same simulations in the corresponding subfigures in Figure 2,3. Other parameters are fixed to: *β_H_* = *β_P_* = 1,*σ* = 0.85. The abbreviations are a follows: S: All genotypes maintained in the given host species have the susceptibility allele at the given locus; R: all genotypes maintained in the given host species have the resistant allele at the respective locus, R/S: genotypes with the resistant allele and genotypes with the susceptible allele at the given locus are maintained in the population. As a reminder, the private locus corresponds to *L*1 in host *M* and *L*2 in host *H* and shared locus corresponds to *L*3 in both host species. By definition, host *H* is always susceptible at locus *L*1 and hence no indication on the status of the maintained alleles at this locus is given. The same applies to host *M* for locus *L*2. For example, the combination private: R, shared: R/S would refer to the maintenance of genotypes 100/101 in host species *M* and 010/011 in host species *H*.

**Figure S5.**
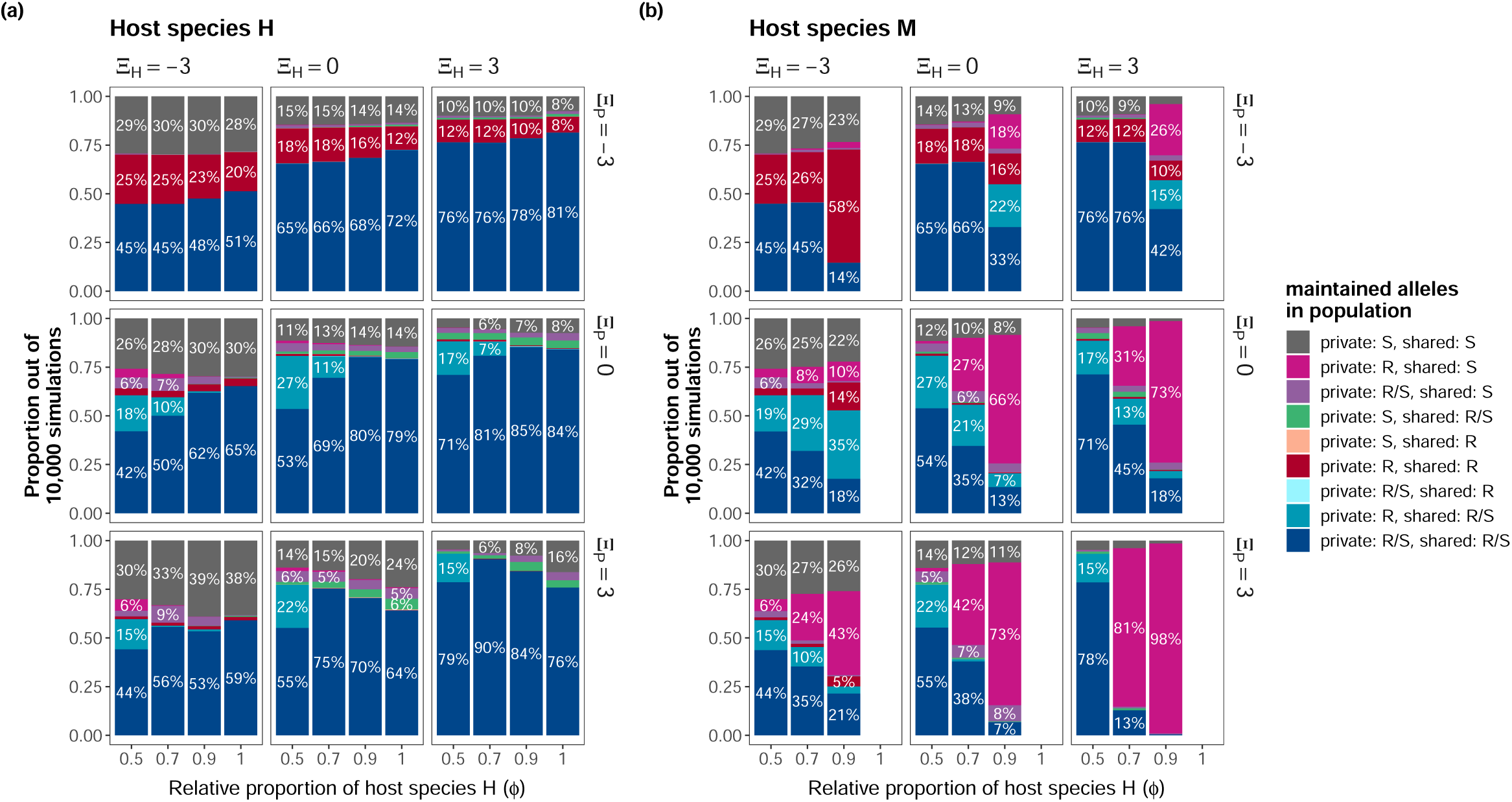
Alleles maintained for (a) host species *H* and (b) host species *M* for different combinations of the shape parameters *Ξ_P_* (rows) and *Ξ_H_* = *Ξ_M_* (columns) for different proportions of host species *H* (*ϕ*) (x-axis). Shown are the aggregated results from 10,000 simulations with the maximum costs of resistance (*Ω_H_*) and virulence (*Ω_P_*) independently drawn from U(0.01, 0.3) and random initialization of genotype frequencies. All combinations that are observed in at least 5% of the analyzed simulations are labeled. Other parameters are fixed to: *β_H_* = *β_P_* = 1,*σ* = 0.85. As a reminder, the private locus corresponds to *L*1 in host *M* and *L*2 in host *H* and shared locus corresponds to *L*3 in both host species. By definition, host *H* is always susceptible at locus *L*1 and hence no indication on the status of the maintained alleles at this locus is given. The same applies to host *M* for locus *L*2. The abbreviations are a follows: S: All genotypes maintained in the given host species have the susceptibility allele at the given locus, R: all genotypes maintained in the given host species have the resistant allele at the respective locus, R/S: both genotypes with the resistant allele and the susceptible allele at the given locus are maintained in the population. For example, the combination private: R, shared: R/S would refer to the maintenance of genotypes 100/101 in host species *M* and 010/011 in host species *H*.

**Figure S6.**
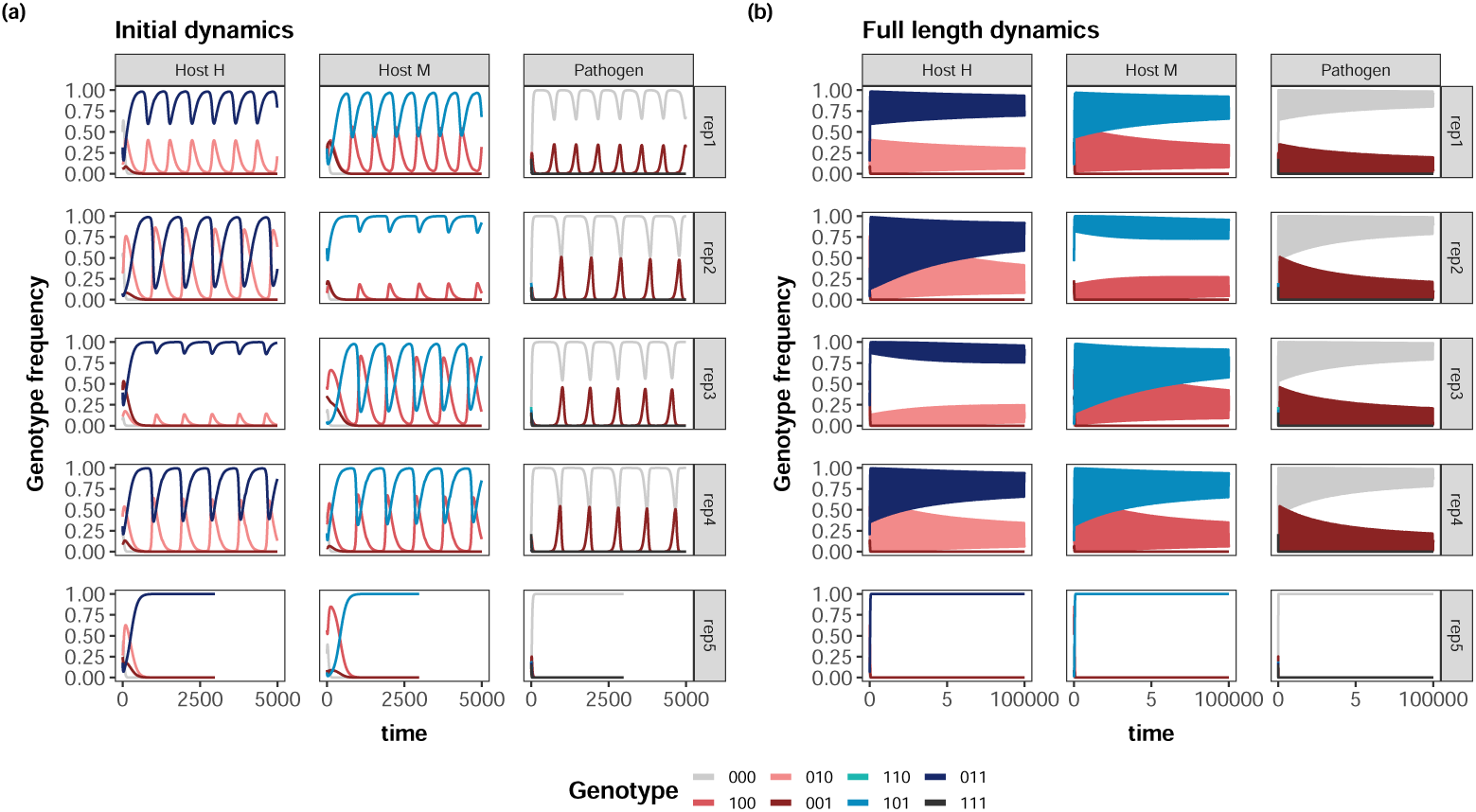
Figure showing the effect of initial genotype frequencies on the maintained genotypes. Shown are the results for 50 simulations for 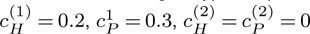 and *ϕ* = 0.5 with random initialization of genotype frequencies, but all other parameters fixed to the same value. Each dot represents the average genotype frequency of a given genotype in a single simulation in the given time interval. Each facet shows the result for a single time interval (columns) for each species (rows).

**Figure S7.**
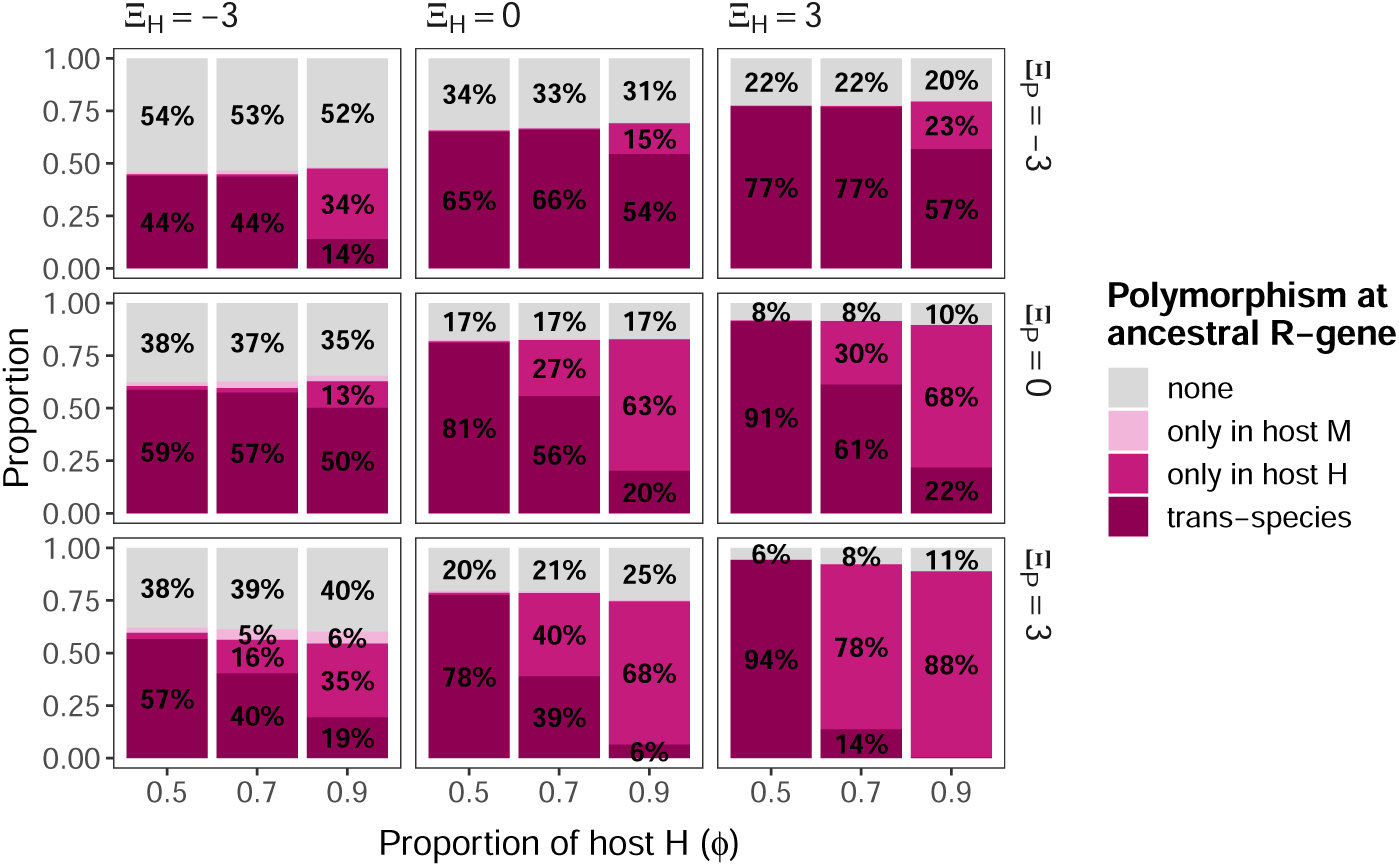
Proportion of simulations where polymorphism at the shared resistance gene is maintained for different shapes of the cost function in the host (columns) and pathogen (rows). Each subplot shows the results for increasing proportions *ϕ* of host species *H*. Each bar presents the results of 10,000 simulations with 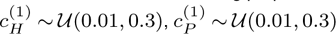 and random initialization of genotype frequencies.

**Figure S8.**
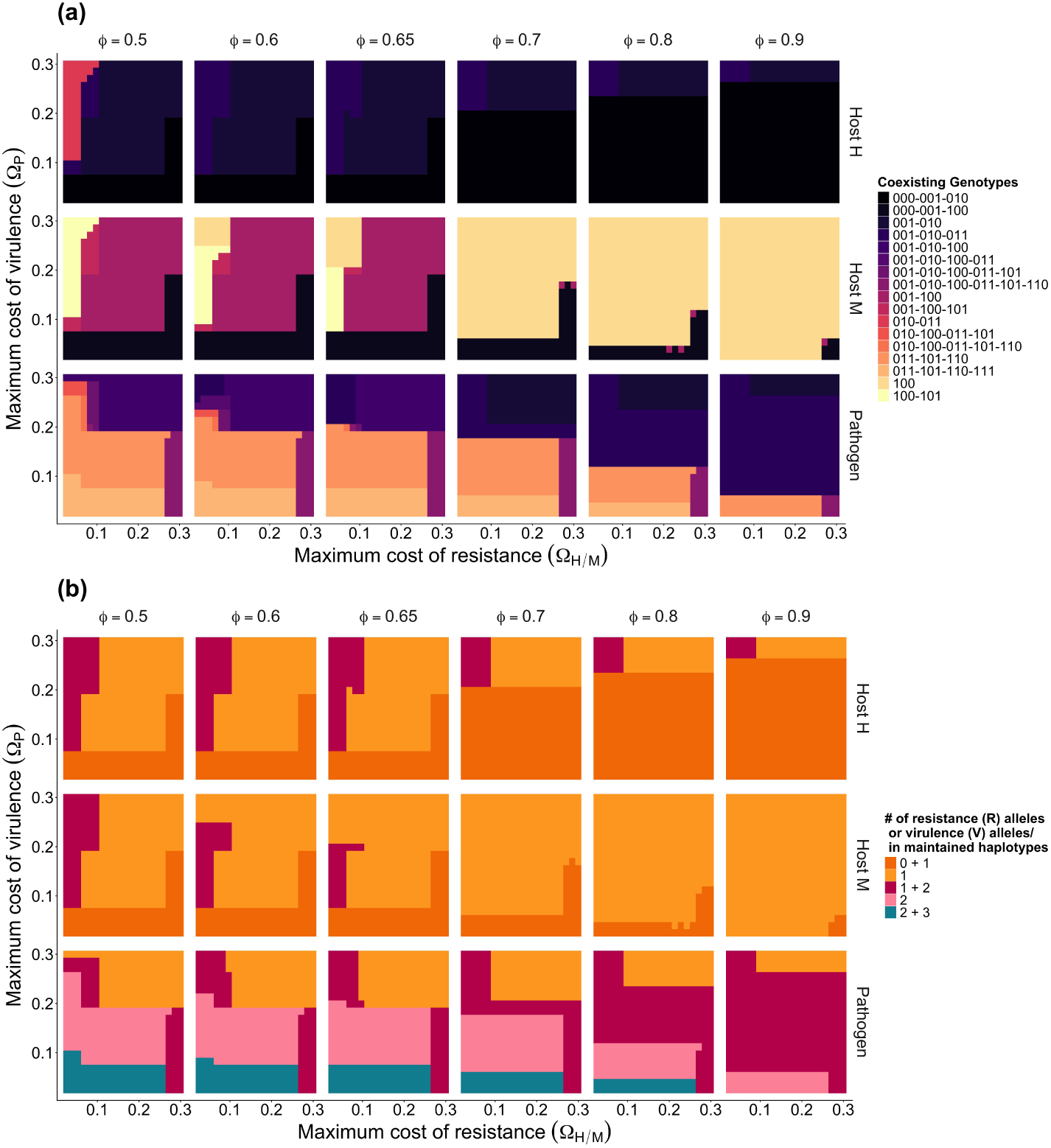
(a) Heatmaps showing the coexisting genotypes across different maximum costs of resistance (*Ω_H_*) and maximum costs of virulence (*Ω_P_*). Columns contain results for different values of the proportion of host *H* while rows show the different species. Colors display which genotypes have a frequency above 10*^−^*^6^. Simulations are conducted over 10^5^ timesteps and results are averaged over the last 10%. Parameters: *β_M_* = *β_H_* = *β_P_* = 1, *Ξ_M_* = *Ξ_H_* = *Ξ_P_* = 3 and *σ* = 0.85. (b) The same data as in (a) but now the colors represent the number of resistance or virulence alleles that coexist in the different states as in Figure 4.

## C. Supplementary Tables

**Table S2.**
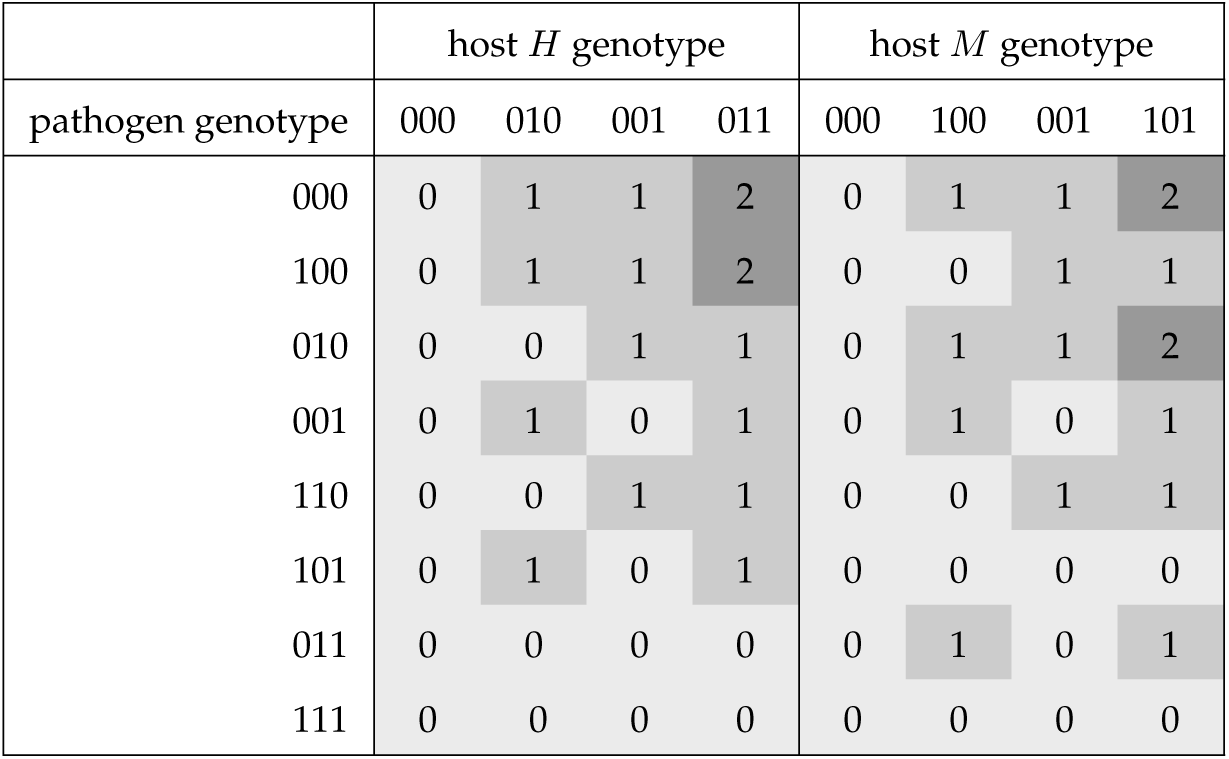
The number of recognizing resistance alleles *d* in all possible pairwise interactions between host genotypes (columns; host *H* left, host *M* right) and pathogen genotypes (rows). The probability of infection in the interactions is calculated as *σ^d^*.

| | host $H$ genotype | | | | host $M$ genotype | | | |
| --- | --- | --- | --- | --- | --- | --- | --- | --- |
| pathogen genotype | 000 | 010 | 001 | 011 | 000 | 100 | 001 | 101 |
| 000 | 0 | 1 | 1 | 2 | 0 | 1 | 1 | 2 |
| 100 | 0 | 1 | 1 | 2 | 0 | 0 | 1 | 1 |
| 010 | 0 | 0 | 1 | 1 | 0 | 1 | 1 | 2 |
| 001 | 0 | 1 | 0 | 1 | 0 | 1 | 0 | 1 |
| 110 | 0 | 0 | 1 | 1 | 0 | 0 | 1 | 1 |
| 101 | 0 | 1 | 0 | 1 | 0 | 0 | 0 | 0 |
| 011 | 0 | 0 | 0 | 0 | 0 | 1 | 0 | 1 |
| 111 | 0 | 0 | 0 | 0 | 0 | 0 | 0 | 0 |

**Table S3.** Detailed overview on the relative proportions underlying Figure 2b.

| Species | $\phi$ | both | ancestral only | private only | none |
| --- | --- | --- | --- | --- | --- |
| Host H | 0.5 | 0.932 | 0.011 | 0.010 x | 0.046 |
| Host M | 0.5 | 0.932 | 0.012 | 0.009 | 0.048 |
| Host H | 0.7 | 0.907 | 0.018 | 0.011 | 0.064 |
| Host M | 0.7 | 0.128 | 0.009 | 0.822 | 0.040 |
| Host H | 0.9 | 0.843 | 0.047 | 0.032 | 0.078 |
| Host M | 0.9 | 0.004 | 0.000 | 0.980 | 0.016 |
| Host H | 1 | 0.758 | 0.038 | 0.041 | 0.163 |
| Host M | 1 | - | - | - | - |

**Table S4.** Detailed overview on the relative proportions underlying Figure 2c.

| Species | $\phi$ | both | ancestral only | private only | none |
| --- | --- | --- | --- | --- | --- |
| Host H | 0.5 | 0.787 | 0.009 | 0.043 | 0.161 |
| Host M | 0.5 | 0.786 | 0.009 | 0.044 | 0.161 |
| Host H | 0.7 | 0.777 | 0.013 | 0.036 | 0.174 |
| Host M | 0.7 | 0.581 | 0.006 | 0.267 | 0.146 |
| Host H | 0.9 | 0.763 | 0.020 | 0.027 | 0.190 |
| Host M | 0.9 | 0.398 | 0.001 | 0.482 | 0.120 |
| Host H | 1 | 0.755 | 0.023 | 0.022 | 0.200 |
| Host M | 1 | - | - | - | - |

**Table S5.** Detailed overview on the relative proportions underlying Figure 3b.

| Species | $\phi$ | both | ancestral only | private only | none |
| --- | --- | --- | --- | --- | --- |
| Host H | 0.5 | 0.785 | 0.158 | 0.009 | 0.047 |
| Host M | 0.5 | 0.784 | 0.159 | 0.008 | 0.049 |
| Host H | 0.7 | 0.904 | 0.018 | 0.011 | 0.067 |
| Host M | 0.7 | 0.128 | 0.009 | 0.009 | 0.854 |
| Host H | 0.9 | 0.842 | 0.047 | 0.032 | 0.079 |
| Host M | 0.9 | 0.004 | 0.000 | 0.005 | 0.991 |
| Host H | 1 | 0.758 | 0.038 | 0.041 | 0.163 |
| Host M | 1 | - | - | - | - |

**Table S6.** Detailed overview on the relative proportions underlying Figure 3c.

| Species | $\phi$ | both | ancestral only | private only | none |
| --- | --- | --- | --- | --- | --- |
| Host H | 0.5 | 0.245 | 0.028 | 0.137 | 0.590 |
| Host M | 0.5 | 0.246 | 0.028 | 0.136 | 0.590 |
| Host H | 0.7 | 0.247 | 0.032 | 0.045 | 0.676 |
| Host M | 0.7 | 0.454 | 0.029 | 0.089 | 0.429 |
| Host H | 0.9 | 0.252 | 0.027 | 0.022 | 0.699 |
| Host M | 0.9 | 0.691 | 0.027 | 0.097 | 0.185 |
| Host H | 1 | 0.253 | 0.022 | 0.022 | 0.703 |
| Host M | 1 | - | - | - | - |

**Table S7.** Detailed summary of the resistance alleles maintained for different combinations of the shape parameters *ΞH*, *ΞP* and relative proportions of host species *H (ϕ)*. Shown are the relative proportions in 10,000 simulations per combination with *ΩH ∼ U*(0.01, 0.3), *ΩP ∼ U*(0.01, 0.3) and random initial genotype frequencies. The results are shown for the exact same simulations used to generate Figure 2c and 3c and 4.

|  |  |  | Host H |  |  |  | Host M |  |  |  |
| --- | --- | --- | --- | --- | --- | --- | --- | --- | --- | --- |
| $\Xi_H$ | $\Xi_P$ | $\phi$ | both | private only | ancestral only | none | both | private only | ancestral only | none |
| -3 | -3 | 0.5 | 0.7003 | 0.0062 | 0 | 0.2935 | 0.7002 | 0.0059 | 1e-04 | 0.2938 |
| -3 | -3 | 0.7 | 0.6993 | 0.0019 | 2e-04 | 0.2986 | 0.7133 | 0.0186 | 1e-04 | 0.268 |
| -3 | -3 | 0.9 | 0.7007 | 0.0023 | 2e-04 | 0.2968 | 0.7263 | 0.0389 | 0 | 0.2348 |
| -3 | -3 | 1 | 0.7139 | 0.0028 | 0.0024 | 0.2809 | 0 | 0 | 0 | 0 |
| -3 | 0 | 0.5 | 0.6411 | 0.1004 | 0 | 0.2585 | 0.6412 | 0.1001 | 0 | 0.2587 |
| -3 | 0 | 0.7 | 0.6276 | 0.0875 | 1e-04 | 0.2848 | 0.6399 | 0.1118 | 0 | 0.2483 |
| -3 | 0 | 0.9 | 0.6615 | 0.0392 | 0.0019 | 0.2974 | 0.6704 | 0.1075 | 0 | 0.2221 |
| -3 | 0 | 1 | 0.6914 | 0.0047 | 0.0042 | 0.2997 | 0 | 0 | 0 | 0 |
| -3 | 3 | 0.5 | 0.6102 | 0.0885 | 0 | 0.3013 | 0.6055 | 0.0934 | 0 | 0.3011 |
| -3 | 3 | 0.7 | 0.5775 | 0.0899 | 3e-04 | 0.3323 | 0.4697 | 0.2562 | 0 | 0.2741 |
| -3 | 3 | 0.9 | 0.5602 | 0.0475 | 0.0012 | 0.3911 | 0.3021 | 0.4377 | 0 | 0.2602 |
| -3 | 3 | 1 | 0.6078 | 0.0033 | 0.0057 | 0.3832 | 0 | 0 | 0 | 0 |
| 0 | -3 | 0.5 | 0.8349 | 0.0173 | 0.0015 | 0.1463 | 0.8326 | 0.0218 | 0.001 | 0.1446 |
| 0 | -3 | 0.7 | 0.8399 | 0.0115 | 0.0021 | 0.1465 | 0.8405 | 0.0295 | 5e-04 | 0.1295 |
| 0 | -3 | 0.9 | 0.8416 | 0.0088 | 0.007 | 0.1426 | 0.7061 | 0.2011 | 3e-04 | 0.0925 |
| 0 | -3 | 1 | 0.8453 | 0.0105 | 0.0091 | 0.1351 | 0 | 0 | 0 | 0 |
| 0 | 0 | 0.5 | 0.8175 | 0.0559 | 0.0119 | 0.1147 | 0.8186 | 0.055 | 0.0101 | 0.1163 |
| 0 | 0 | 0.7 | 0.8166 | 0.038 | 0.0191 | 0.1263 | 0.5658 | 0.3299 | 0.0051 | 0.0992 |
| 0 | 0 | 0.9 | 0.8081 | 0.0302 | 0.0252 | 0.1365 | 0.2092 | 0.706 | 4e-04 | 0.0844 |
| 0 | 0 | 1 | 0.795 | 0.0282 | 0.0322 | 0.1446 | 0 | 0 | 0 | 0 |
| 0 | 3 | 0.5 | 0.7724 | 0.0729 | 0.015 | 0.1397 | 0.773 | 0.0727 | 0.0135 | 0.1408 |
| 0 | 3 | 0.7 | 0.7543 | 0.0582 | 0.034 | 0.1535 | 0.3903 | 0.4808 | 0.0078 | 0.1211 |
| 0 | 3 | 0.9 | 0.7053 | 0.0528 | 0.0435 | 0.1984 | 0.0686 | 0.8152 | 0.0033 | 0.1129 |
| 0 | 3 | 1 | 0.6413 | 0.0588 | 0.0599 | 0.24 | 0 | 0 | 0 | 0 |
| 3 | -3 | 0.5 | 0.8803 | 0.0107 | 0.0096 | 0.0994 | 0.8806 | 0.011 | 0.0099 | 0.0985 |
| 3 | -3 | 0.7 | 0.8809 | 0.0067 | 0.0103 | 0.1021 | 0.8837 | 0.0185 | 0.0061 | 0.0917 |
| 3 | -3 | 0.9 | 0.8846 | 0.0076 | 0.0089 | 0.0989 | 0.6696 | 0.2906 | 3e-04 | 0.0395 |
| 3 | -3 | 1 | 0.8957 | 0.0144 | 0.0137 | 0.0762 | 0 | 0 | 0 | 0 |
| 3 | 0 | 0.5 | 0.8919 | 0.0262 | 0.0337 | 0.0482 | 0.8944 | 0.0276 | 0.0305 | 0.0475 |
| 3 | 0 | 0.7 | 0.8906 | 0.0198 | 0.034 | 0.0556 | 0.5963 | 0.336 | 0.0273 | 0.0404 |
| 3 | 0 | 0.9 | 0.8614 | 0.024 | 0.0409 | 0.0737 | 0.2225 | 0.7622 | 0.0013 | 0.014 |
| 3 | 0 | 1 | 0.8475 | 0.0388 | 0.0386 | 0.0751 | 0 | 0 | 0 | 0 |
| 3 | 3 | 0.5 | 0.9325 | 0.0101 | 0.0109 | 0.0465 | 0.9317 | 0.0087 | 0.0116 | 0.048 |
| 3 | 3 | 0.7 | 0.9068 | 0.0113 | 0.0175 | 0.0644 | 0.1285 | 0.8223 | 0.0093 | 0.0399 |
| 3 | 3 | 0.9 | 0.8429 | 0.0321 | 0.0472 | 0.0778 | 0.0038 | 0.9804 | 1e-04 | 0.0157 |
| 3 | 3 | 1 | 0.758 | 0.041 | 0.038 | 0.163 | 0 | 0 | 0 | 0 |

**Table S8.** Detailed summary of the R gene polymorphisms maintained for different combinations of the shape parameters *ΞH*, *ΞP* and relative proportions of host species *H (ϕ)*. Shown are the relative proportions in 10,000 simulations per combination with *ΩH ∼ U*(0.01, 0.3), *ΩP ∼ U*(0.01, 0.3) and random initial genotype frequencies. The results are shown for the exact same simulations used to generate Figure 2c and 3c and 4.

|  |  |  | Host H |  |  |  | Host M |  |  |  |
| --- | --- | --- | --- | --- | --- | --- | --- | --- | --- | --- |
| $\Xi_H$ | $\Xi_P$ | $\phi$ | both | private only | ancestral only | none | both | private only | ancestral only | none |
| -3 | -3 | 0.5 | 0.4474 | 0.0056 | 2e-04 | 0.5468 | 0.4478 | 0.0048 | 3e-04 | 0.5471 |
| -3 | -3 | 0.7 | 0.4476 | 0.0017 | 1e-04 | 0.5506 | 0.4545 | 0.0124 | 1e-04 | 0.533 |
| -3 | -3 | 0.9 | 0.4751 | 0.002 | 1e-04 | 0.5228 | 0.1447 | 0.0086 | 7e-04 | 0.846 |
| -3 | -3 | 1 | 0.5126 | 0.0025 | 0.0017 | 0.4832 | 0 | 0 | 0 | 0 |
| -3 | 0 | 0.5 | 0.4199 | 0.0554 | 0.1849 | 0.3398 | 0.4191 | 0.0559 | 0.1861 | 0.3389 |
| -3 | 0 | 0.7 | 0.4997 | 0.0655 | 0.0953 | 0.3395 | 0.3196 | 0.0279 | 0.2865 | 0.366 |
| -3 | 0 | 0.9 | 0.6173 | 0.0376 | 0.0102 | 0.3349 | 0.1764 | 0.0081 | 0.3513 | 0.4642 |
| -3 | 0 | 1 | 0.6526 | 0.0047 | 0.0042 | 0.3385 | 0 | 0 | 0 | 0 |
| -3 | 3 | 0.5 | 0.4412 | 0.0287 | 0.1549 | 0.3752 | 0.4369 | 0.0322 | 0.1541 | 0.3768 |
| -3 | 3 | 0.7 | 0.5558 | 0.0863 | 0.0073 | 0.3506 | 0.3532 | 0.0169 | 0.0999 | 0.53 |
| -3 | 3 | 0.9 | 0.5338 | 0.0469 | 0.0112 | 0.4081 | 0.2135 | 0.0067 | 0.0367 | 0.7431 |
| -3 | 3 | 1 | 0.5904 | 0.0033 | 0.0057 | 0.4006 | 0 | 0 | 0 | 0 |
| 0 | -3 | 0.5 | 0.6536 | 0.0156 | 0.0025 | 0.3283 | 0.6517 | 0.0198 | 0.002 | 0.3265 |
| 0 | -3 | 0.7 | 0.6623 | 0.01 | 0.0029 | 0.3248 | 0.6603 | 0.0247 | 0.003 | 0.312 |
| 0 | -3 | 0.9 | 0.6831 | 0.0081 | 0.0067 | 0.3021 | 0.3286 | 0.0248 | 0.2195 | 0.4271 |
| 0 | -3 | 1 | 0.7243 | 0.0098 | 0.0084 | 0.2575 | 0 | 0 | 0 | 0 |
| 0 | 0 | 0.5 | 0.5347 | 0.044 | 0.2838 | 0.1375 | 0.5387 | 0.0433 | 0.279 | 0.139 |
| 0 | 0 | 0.7 | 0.6948 | 0.0325 | 0.1292 | 0.1435 | 0.3454 | 0.0553 | 0.2157 | 0.3836 |
| 0 | 0 | 0.9 | 0.8009 | 0.0301 | 0.0267 | 0.1423 | 0.1342 | 0.0459 | 0.0705 | 0.7494 |
| 0 | 0 | 1 | 0.7915 | 0.0285 | 0.0322 | 0.1478 | 0 | 0 | 0 | 0 |
| 0 | 3 | 0.5 | 0.551 | 0.0561 | 0.2352 | 0.1577 | 0.5523 | 0.0549 | 0.2329 | 0.1599 |
| 0 | 3 | 0.7 | 0.7535 | 0.0508 | 0.032 | 0.1637 | 0.3788 | 0.0652 | 0.0186 | 0.5374 |
| 0 | 3 | 0.9 | 0.705 | 0.0483 | 0.0405 | 0.2062 | 0.0673 | 0.0823 | 0.0036 | 0.8468 |
| 0 | 3 | 1 | 0.6411 | 0.054 | 0.0554 | 0.2495 | 0 | 0 | 0 | 0 |
| 3 | -3 | 0.5 | 0.7628 | 0.0107 | 0.0103 | 0.2162 | 0.7637 | 0.0111 | 0.0103 | 0.2149 |
| 3 | -3 | 0.7 | 0.7615 | 0.0067 | 0.011 | 0.2208 | 0.7626 | 0.0163 | 0.0077 | 0.2134 |
| 3 | -3 | 0.9 | 0.7842 | 0.008 | 0.0092 | 0.1986 | 0.421 | 0.027 | 0.1481 | 0.4039 |
| 3 | -3 | 1 | 0.8133 | 0.0145 | 0.0138 | 0.1584 | 0 | 0 | 0 | 0 |
| 3 | 0 | 0.5 | 0.7098 | 0.0249 | 0.2063 | 0.059 | 0.7122 | 0.0262 | 0.2036 | 0.058 |
| 3 | 0 | 0.7 | 0.8083 | 0.0197 | 0.1058 | 0.0662 | 0.4544 | 0.0307 | 0.1601 | 0.3548 |
| 3 | 0 | 0.9 | 0.8523 | 0.0263 | 0.0435 | 0.0779 | 0.1792 | 0.0354 | 0.0384 | 0.747 |
| 3 | 0 | 1 | 0.8409 | 0.0407 | 0.0399 | 0.0785 | 0 | 0 | 0 | 0 |
| 3 | 3 | 0.5 | 0.7854 | 0.0093 | 0.1579 | 0.0474 | 0.7845 | 0.0079 | 0.1587 | 0.0489 |
| 3 | 3 | 0.7 | 0.9038 | 0.0113 | 0.0175 | 0.0674 | 0.1284 | 0.0087 | 0.0094 | 0.8535 |
| 3 | 3 | 0.9 | 0.8415 | 0.0321 | 0.0472 | 0.0792 | 0.0038 | 0.0052 | 1e-04 | 0.9909 |
| 3 | 3 | 1 | 0.7579 | 0.041 | 0.038 | 0.1631 | 0 | 0 | 0 | 0 |

**Table S9.** Detailed summary of the genotype combinations maintained in host *H* for different combinations of the shape parameters *Ξ_H_*, *Ξ_P_* and relative proportions of host species *H* (*ϕ*). Shown are the relative proportions in 10,000 simulations per combination with *Ω_H_* ∼ *U*(0.01, 0.3), Ω_P_ ∼ U(0.01, 0.3) and random initial genotype frequencies. The results are shown for the exact same simulations used to generate Figure 2c, 3c and 4.

| $\Xi_H$ | $\Xi_P$ | $\phi$ | 0 | 1 | 0 + 1 | 2 | 0 + 2 | 1 + 2 | 0 + 1 + 2 |
| --- | --- | --- | --- | --- | --- | --- | --- | --- | --- |
| -3 | -3 | 0.5 | 0.2935 | 0.0085 | 0.0056 | 0.2527 | 0.4389 | 8e-04 | 0 |
| -3 | -3 | 0.7 | 0.2986 | 0.0068 | 0.0016 | 0.2515 | 0.4408 | 7e-04 | 0 |
| -3 | -3 | 0.9 | 0.2968 | 0.0086 | 0.0021 | 0.2255 | 0.4664 | 6e-04 | 0 |
| -3 | -3 | 1.0 | 0.2809 | 0.0106 | 0.0045 | 0.201 | 0.5022 | 8e-04 | 0 |
| -3 | 0 | 0.5 | 0.2585 | 0.045 | 0.0554 | 0.0363 | 0.2113 | 0.1849 | 0.2086 |
| -3 | 0 | 0.7 | 0.2848 | 0.0225 | 0.0656 | 0.0327 | 0.1957 | 0.0952 | 0.3035 |
| -3 | 0 | 0.9 | 0.2974 | 0.0037 | 0.0395 | 0.0359 | 0.2761 | 0.0083 | 0.3391 |
| -3 | 0 | 1.0 | 0.2997 | 8e-04 | 0.009 | 0.0388 | 0.6517 | 0 | 0 |
| -3 | 3 | 0.5 | 0.3013 | 0.0598 | 0.0287 | 0.0141 | 0.1964 | 0.1549 | 0.2448 |
| -3 | 3 | 0.7 | 0.3323 | 0.0036 | 0.0866 | 0.0147 | 0.3259 | 0.007 | 0.2299 |
| -3 | 3 | 0.9 | 0.3911 | 6e-04 | 0.0481 | 0.0164 | 0.5039 | 0.01 | 0.0299 |
| -3 | 3 | 1.0 | 0.3832 | 0 | 0.009 | 0.0174 | 0.5904 | 0 | 0 |
| 0 | -3 | 0.5 | 0.1463 | 0.1763 | 0.3043 | 0.1795 | 0 | 0.1936 | 0 |
| 0 | -3 | 0.7 | 0.1465 | 0.1752 | 0.3022 | 0.1763 | 0 | 0.1998 | 0 |
| 0 | -3 | 0.9 | 0.1426 | 0.1727 | 0.3046 | 0.1573 | 0 | 0.2228 | 0 |
| 0 | -3 | 1.0 | 0.1351 | 0.1654 | 0.354 | 0.1201 | 0 | 0.2254 | 0 |
| 0 | 0 | 0.5 | 0.1147 | 0.1683 | 0.3427 | 0.0103 | 0 | 0.3625 | 0.0015 |
| 0 | 0 | 0.7 | 0.1263 | 0.1637 | 0.3576 | 0.0098 | 3e-04 | 0.3375 | 0.0048 |
| 0 | 0 | 0.9 | 0.1365 | 0.1481 | 0.4558 | 0.0031 | 0 | 0.2506 | 0.0059 |
| 0 | 0 | 1.0 | 0.1446 | 0.1327 | 0.5162 | 0.0014 | 0 | 0.1991 | 0.006 |
| 0 | 3 | 0.5 | 0.1397 | 0.1052 | 0.458 | 2e-04 | 0 | 0.2939 | 0.003 |
| 0 | 3 | 0.7 | 0.1535 | 0.0558 | 0.6592 | 8e-04 | 0 | 0.1205 | 0.0102 |
| 0 | 3 | 0.9 | 0.1984 | 0.0565 | 0.6857 | 3e-04 | 0 | 0.0515 | 0.0076 |
| 0 | 3 | 1.0 | 0.24 | 0.0633 | 0.6781 | 2e-04 | 0 | 0.0134 | 0.005 |
| 3 | -3 | 0.5 | 0.0994 | 0.3208 | 0.3313 | 0.1167 | 0 | 0.1318 | 0 |
| 3 | -3 | 0.7 | 0.1021 | 0.3118 | 0.3356 | 0.1186 | 0 | 0.1319 | 0 |
| 3 | -3 | 0.9 | 0.0989 | 0.3095 | 0.343 | 0.0996 | 0 | 0.149 | 0 |
| 3 | -3 | 1.0 | 0.0762 | 0.2985 | 0.4032 | 0.0821 | 0 | 0.14 | 0 |
| 3 | 0 | 0.5 | 0.0482 | 0.3473 | 0.3657 | 0.0094 | 0 | 0.2294 | 0 |
| 3 | 0 | 0.7 | 0.0556 | 0.3153 | 0.401 | 0.0096 | 0 | 0.2185 | 0 |
| 3 | 0 | 0.9 | 0.0737 | 0.2598 | 0.5157 | 0.0042 | 0 | 0.1466 | 0 |
| 3 | 0 | 1.0 | 0.0751 | 0.2306 | 0.5704 | 0.0034 | 0 | 0.1205 | 0 |
| 3 | 3 | 0.5 | 0.0465 | 0.5636 | 0.1991 | 1e-04 | 0 | 0.1907 | 0 |
| 3 | 3 | 0.7 | 0.0644 | 0.2779 | 0.5966 | 0.003 | 0 | 0.0581 | 0 |
| 3 | 3 | 0.9 | 0.0778 | 0.1322 | 0.765 | 0.0014 | 0 | 0.0236 | 0 |
| 3 | 3 | 1.0 | 0.163 | 0.0694 | 0.7649 | 1e-04 | 0 | 0.0026 | 0 |

**Table S10.** Detailed summary of the genotype combinations maintained in host *M* for different combinations of the shape parameters *Ξ_H_*, *Ξ_P_* and relative proportions of host species *H* (*ϕ*). Shown are the relative proportions in 10,000 simulations per combination with *Ω_H_* ∼ U(0.01, 0.3), *Ω_P_* ∼ U(0.01, 0.3) and random initial genotype frequencies. The results are shown for the exact same simulations used to generate Figure 2c, 3c and 4.

| $\Xi_H$ | $\Xi_P$ | $\phi$ | 0 | 1 | 0 + 1 | 2 | 0 + 2 | 1 + 2 | 0 + 1 + 2 |
| --- | --- | --- | --- | --- | --- | --- | --- | --- | --- |
| -3 | -3 | 0.5 | 0.2938 | 0.0089 | 0.0049 | 0.2521 | 0.4396 | 7e-04 | 0 |
| -3 | -3 | 0.7 | 0.268 | 0.0112 | 0.0123 | 0.2586 | 0.4494 | 5e-04 | 0 |
| -3 | -3 | 0.9 | 0.2348 | 0.0303 | 0.0086 | 0.5809 | 0.1447 | 7e-04 | 0 |
| -3 | 0 | 0.5 | 0.2587 | 0.0444 | 0.0559 | 0.036 | 0.2114 | 0.1861 | 0.2075 |
| -3 | 0 | 0.7 | 0.2483 | 0.0839 | 0.0279 | 0.0338 | 0.2153 | 0.2865 | 0.1043 |
| -3 | 0 | 0.9 | 0.2221 | 0.0994 | 0.0081 | 0.1427 | 0.1741 | 0.3513 | 0.0023 |
| -3 | 3 | 0.5 | 0.3011 | 0.0612 | 0.0322 | 0.0145 | 0.1918 | 0.1541 | 0.2451 |
| -3 | 3 | 0.7 | 0.2741 | 0.2393 | 0.0169 | 0.0166 | 0.2154 | 0.0999 | 0.1378 |
| -3 | 3 | 0.9 | 0.2602 | 0.431 | 0.0067 | 0.0519 | 0.2044 | 0.0367 | 0.0091 |
| 0 | -3 | 0.5 | 0.1446 | 0.1764 | 0.3066 | 0.1789 | 0 | 0.1935 | 0 |
| 0 | -3 | 0.7 | 0.1295 | 0.1784 | 0.3153 | 0.1774 | 0 | 0.1994 | 0 |
| 0 | -3 | 0.9 | 0.0925 | 0.3535 | 0.1751 | 0.1582 | 0 | 0.2207 | 0 |
| 0 | 0 | 0.5 | 0.1163 | 0.1679 | 0.3413 | 0.0102 | 0 | 0.3625 | 0.0018 |
| 0 | 0 | 0.7 | 0.0992 | 0.4158 | 0.2636 | 0.0094 | 0 | 0.2113 | 7e-04 |
| 0 | 0 | 0.9 | 0.0844 | 0.7848 | 0.0556 | 0.0048 | 1e-04 | 0.0703 | 0 |
| 0 | 3 | 0.5 | 0.1408 | 0.1064 | 0.457 | 0 | 0 | 0.2932 | 0.0026 |
| 0 | 3 | 0.7 | 0.1211 | 0.4523 | 0.4147 | 0 | 0 | 0.0115 | 4e-04 |
| 0 | 3 | 0.9 | 0.1129 | 0.7664 | 0.1194 | 7e-04 | 0 | 6e-04 | 0 |
| 3 | -3 | 0.5 | 0.0985 | 0.321 | 0.3321 | 0.1161 | 0 | 0.1323 | 0 |
| 3 | -3 | 0.7 | 0.0917 | 0.317 | 0.3407 | 0.1194 | 0 | 0.1312 | 0 |
| 3 | -3 | 0.9 | 0.0395 | 0.5636 | 0.1483 | 0.1007 | 0 | 0.1479 | 0 |
| 3 | 0 | 0.5 | 0.0475 | 0.3484 | 0.3659 | 0.0089 | 0 | 0.2293 | 0 |
| 3 | 0 | 0.7 | 0.0404 | 0.5456 | 0.2721 | 0.0091 | 0 | 0.1328 | 0 |
| 3 | 0 | 0.9 | 0.014 | 0.8958 | 0.0469 | 0.0062 | 0 | 0.0371 | 0 |
| 3 | 3 | 0.5 | 0.048 | 0.5635 | 0.1976 | 1e-04 | 0 | 0.1908 | 0 |
| 3 | 3 | 0.7 | 0.0399 | 0.817 | 0.143 | 0 | 0 | 1e-04 | 0 |
| 3 | 3 | 0.9 | 0.0157 | 0.9756 | 0.0087 | 0 | 0 | 0 | 0 |

**Table S11.**
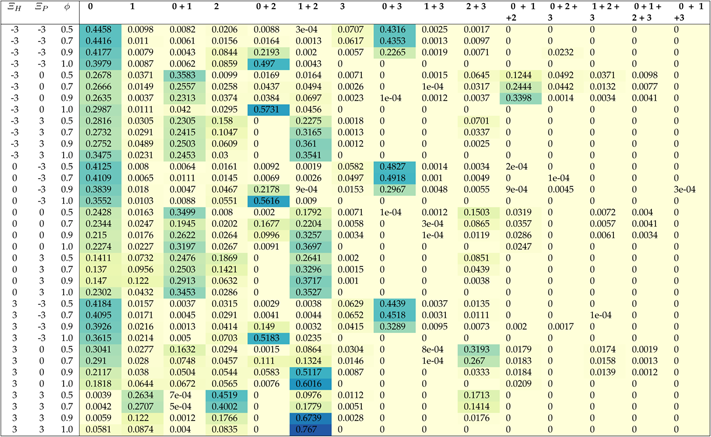
Detailed summary of the genotype combinations maintained in the pathogen *P* for different combinations of the shape parameters *ΞH*, *ΞP* and relative proportions of host species *H* (*ϕ*). Shown are the relative proportions in 10,000 simulations per combination with *ΩH* ∼ U(0.01, 0.3), *ΩP* ∼ U(0.01, 0.3) and random initial genotype frequencies. The results are shown for the exact same simulations used to generate Figure 2c, 3c and 4.

